# TCGA_DEPMAP_ – Mapping Translational Dependencies and Synthetic Lethalities within The Cancer Genome Atlas

**DOI:** 10.1101/2022.03.24.485544

**Authors:** Xu Shi, Christos Gekas, Daniel Verduzco, Sakina Petiwala, Cynthia Jeffries, Charles Lu, Erin Murphy, Tifani Anton, Andy H. Vo, Zoe Xiao, Padmini Narayanan, J. Matthew Barnes, Somdutta Roy, Cyril Ramathal, Michael J. Flister, Zoltan Dezso

## Abstract

The Cancer Genome Atlas (TCGA) has yielded unprecedented genetic and molecular characterization of the cancer genome, yet the functional consequences and patient-relevance of many putative cancer drivers remain undefined. TCGA_DEPMAP_ is the first hybrid map of translational tumor dependencies that was built from machine learning of gene essentiality in the Cancer Dependency Map (DEPMAP) and then translated to TCGA patients. TCGA_DEPMAP_ captured well-known and novel cancer lineage dependencies, oncogenes, and synthetic lethalities, demonstrating the robustness of TCGA_DEPMAP_ as a translational dependency map. Exploratory analyses of TCGA_DEPMAP_ also unveiled novel synthetic lethalities, including the dependency of *PAPSS1* driven by loss of *PAPSS2* which is collaterally deleted with the tumor suppressor gene *PTEN*. Synthetic lethality of *PAPSS1/2* was validated in vitro and in vivo, including the underlying mechanism of synthetic lethality caused by the loss of protein sulfonation that requires *PAPSS1* or *PAPSS2*. Moreover, TCGA_DEPMAP_ demonstrated that patients with predicted *PAPSS1/2* synthetic lethality have worse overall survival, suggesting that these patients are in greater need of drug discovery efforts to target *PAPSS1*. Other map “extensions” were built to capture unique aspects of patient-relevant tumor dependencies using the flexible analytical framework of TCGA_DEPMAP_, including translating gene essentiality to drug response in patient-derived xenograft (PDX) models (i.e., PDXE_DEPMAP_) and predicting gene tolerability within normal tissues (GTEX_DEPMAP_). Collectively, this study demonstrates how translational dependency maps can be used to leverage the rapidly expanding catalog of patient genomic datasets to identify and prioritize novel therapeutic targets with the best therapeutic indices.

## INTRODUCTION

The rapid expansion of genomic technologies to characterize healthy and diseased patient populations has provided unprecedented resolution to the pathophysiological drivers of cancer and many other diseases. In 2018, TCGA completed a 10-year study of 33 tumor types across ∼11,000 patients, which has broadly illuminated the genetic underpinnings of cancer (*1*). Building on the success of TCGA, multiple other initiatives have been launched to explore aspects of cancer initiation, evolution, metastasis, and response to therapy (*2*–*6*), with the hope that the deepening molecular characterization of cancer will improve diagnosis, treatment, and prevention. However, a critical step towards fully leveraging patient data to eradicate cancer is to assign functionality to the observations made in TCGA that translate putative tumor dependencies to life-saving therapies.

One approach to understanding tumor dependencies is through genomewide genetic and chemical perturbation datasets (e.g., DEPMAP (*7, 8*), Project SCORE (*9*), and Connectivity Map (*10*)) that have been paired with thousands of deeply characterized cancer models (e.g., Cancer Cell Line Encyclopedia (*11*), Cancer Cell Line Factory, (*12*), and Human Cancer Models Initiative (*13*)). Multiple studies have demonstrated the ability of DEPMAP to translate gene essentially to novel therapeutic targets (*14*–*18*) and a broader functional understanding of tumor dependencies (*19, 20*). Compared with TCGA, a differentiating strength of the “dependency maps” is that hypotheses can be readily tested, replicated, and refined in different contexts, whereas patient datasets are typically not amenable to functional experimentation. However, the dependency maps also pose limitations when compared to the translatability of TCGA, as homogeneous cell lines in culture dishes do not replicate the pathophysiological complexities of the intact tumor microenvironment (TME) (*21*). Further, the current experimental models do not completely recapitulate the genetic drivers that are present in the patient population (*22*), and experimental outcomes of genetic perturbation screens do not capture most aspects of disease outcome and patient survival.

To address the unique challenges posed by TCGA and DEPMAP, a hybrid dependency map (TCGA_DEPMAP_) was built by machine learning of gene essentiality in the cell based DEPMAP that was translated to TCGA patients. As such, TCGA_DEPMAP_ leverages the experimental strengths of DEPMAP, while enabling patient-relevant translatability of TCGA. TCGA_DEPMAP_ captured well-known cancer lineage dependencies, oncogenes, and synthetic lethalities, demonstrating the robustness of TCGA_DEPMAP_ as a translational dependency map. Exploratory analyses using TCGA_DEPMAP_ also revealed novel tumor dependencies and synthetic lethalities, which could be translated to treatment response and clinical outcomes. Finally, the flexible framework of TCGA_DEPMAP_ enabled the assembly of map “extensions” that captured other aspects of patient-relevant tumor dependencies, including translating dependencies to drug responses in PDX models (i.e., PDXE_DEPMAP_) and predicting gene essentiality within normal tissues (GTEX_DEPMAP_). Collectively, this study demonstrates how translational dependency maps can be leveraged to functionalize oncogenic drivers and novel therapeutic targets in the rapidly expanding catalog of patient genomic datasets.

## RESULTS

### Predictive modelling of gene essentiality

To begin building translational dependency maps, predictive models of gene essentiality were generated from the DEPMAP (*7*) using elastic-net regularization for feature selection and modeling (*23*) (**Figure 1A**). Genomewide gene effect scores for DEPMAP cancer cell models (n = 897) were estimated by CERES (*24*), which measures the dependency probability of each gene relative to the distribution of effect sizes for common essential and nonessential genes within each cell line (*25*). Because many genes do not impact cell viability, elastic-net models were attempted only for genes with at least five dependent and non-dependent cell lines, which included 7,260 out of 18,119 genes (40%) with effects scores in the DEPMAP. In addition to gene effect scores, the input variables for elastic-net predictive modelling included genome-wide gene expression, mutation, and copy number profiles for each cancer cell model. Based on prior evidence that predictive modeling of gene essentiality with RNA expression outperformed similar modeling with DNA features (*26, 27*), two sets of elastic-net models were generated with RNA alone (i.e., expression-only) or combined with mutation and copy number profiles (i.e., multi-omics). Finally, the best fitting elastic-net models were selected by a 10-fold cross-validation to identify models with the minimum error, while balancing the predictive performance with the number of features selected (see Methods for details).

**Figure 1.**
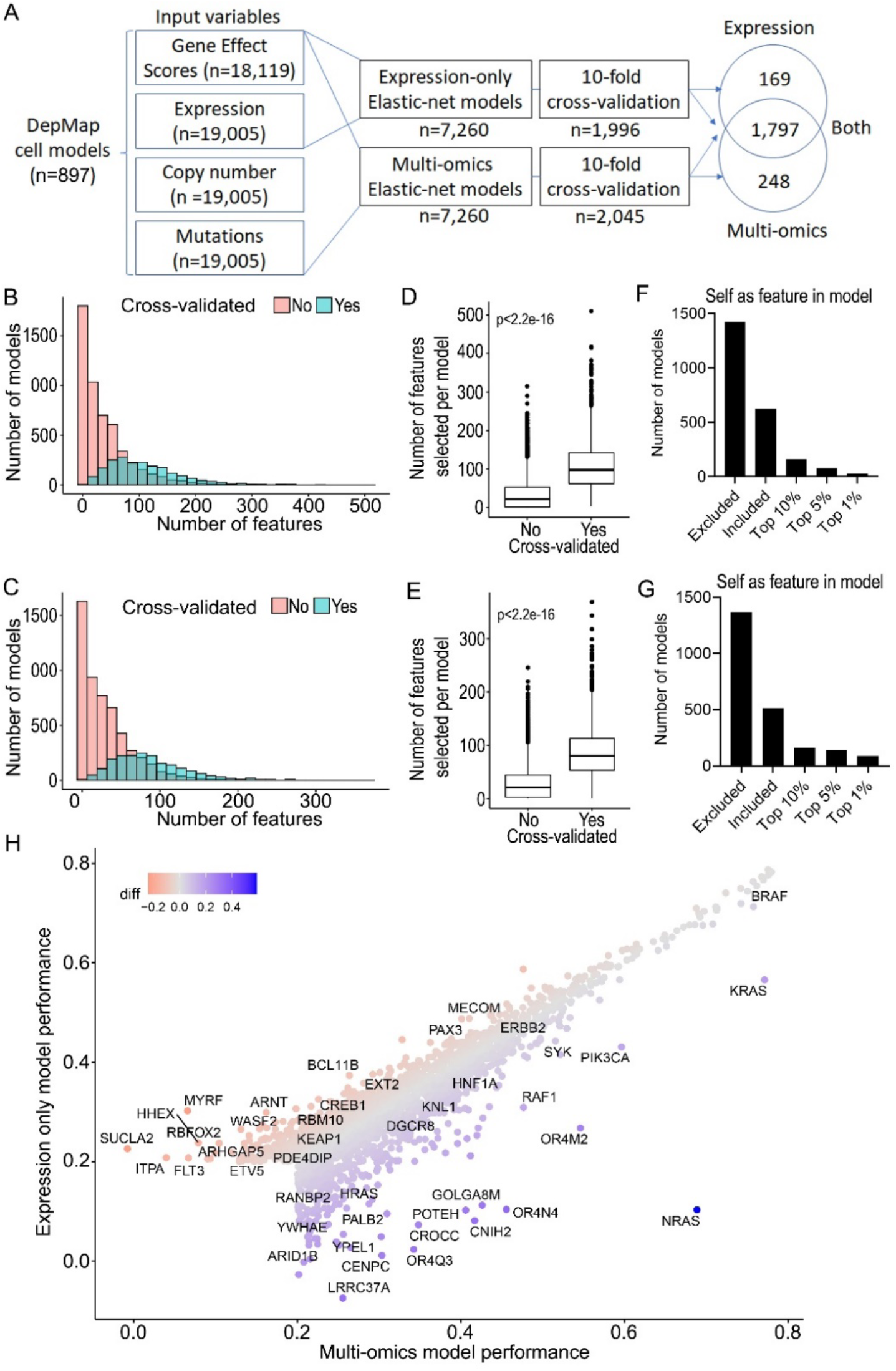
Predictive modeling of gene essentiality in the DEPMAP. (**A**) Schematic of elastic-net models for predictive modeling of gene essentiality in the DEPMAP using expression-only data or multi-omics data. Note the broad overlap in cross-validated models using expression-only or multi-omics data. (**B**) Distribution of features per multi-omics models. (**C**) Number of features per multi-omics model that passed or failed cross-validation. (**D**) Distribution of the target gene (i.e., self) as a feature in the cross-validated multi-omics models. (**E**) Distribution of features per expression-only models. (**F**) Number of features per expression-only model that passed or failed cross-validation. (**G**) Distribution of the target gene (i.e., self) as a feature in the cross-validated expression-only models. (**H**) Comparison of model performance (correlation coefficients) of cross-validated models from multi-omics and expression-only data. Note for (**B-H**) that the performance and characteristics of multi-omics and expression-only models are very similar. All p-values indicated on graphs, as determined by the Wilcoxon rank sum test for two group comparison and Kruskal-Wallis followed by Wilcoxon rank sum test with multiple test correction for the multi-group comparison

The elastic-net models for predicting essentiality of the 7,260 genes (as described above) were compared by 10-fold cross-validation (Pearson’s *r* > 0.2; FDR < 1e-3) when considering expression-only or multi-omics data as input variables (**Tables S1, S2**). The distribution of features per model skewed higher in the multi-omics models (3 to 510 features, median = 98) (**Figure 1B**) compared with the expression-only models (3 to 369 features, median = 80) (**Figure 1C**), and the performance of both improved with the number of features per model (**Figure 1D, E**). Of the 7,260 models, cross-validation confirmed 1,996 expression-only models and 2,045 multi-omics models, of which most cross-validated models overlapped (n=1,797) between the two datasets (**Table S3**). The incidence of self-inclusion of the target gene in the cross-validated models was also similar between multi-omics dataset (31% of models) (**Figure 1F**) and expression-only dataset (26% of models) (**Figure 1G**). Finally, the predictive accuracy of most cross-validated expression-only and multi-omics models performed comparably (e.g., *HER2, BRAF, PIK3CA*, etc.), with a few notable examples that included the oncogenes: *NRAS, FLT3*, and *ARNT* (**Figure 1H**). Collectively, these data demonstrate that predictive models of gene essentiality with expression-only and multi-omics data as input variables perform equivalently in detecting the selective vulnerabilities of cancer.

### Building a translational resource for tumor dependencies: TCGA_DEPMAP_

The schematic in **Figure 2A** outlines the approach to transposing the cross-validated gene essentiality models from the cell-based DEPMAP onto TCGA patients (n = 9,596) to build a translational dependency map (i.e., TCGA_DEPMAP_). TCGA_DEPMAP_ was built using the expression-only elastic-net models of gene essentiality, based on the evidence here (**Figure 1**) and elsewhere (*26, 27*) that only marginal gains were made in model performance by including genomic features. Secondly, transcriptomics data are broadly captured across TCGA and many other clinical studies (*28*), as well as PDX studies (*29, 30*), whereas genomic data are less frequently captured and have greater discrepancies in representation across cancer models and patients. Thirdly, because genetic information is withheld from the expression-only elastic-net models, the transposed essentiality scores can be correlated with genetic drivers in TCGA_DEPMAP_ patients that might otherwise be missed in cancer cell models. Finally, expression-based predictive modeling of dependency can be extended to non-oncological studies (e.g., GTEX), which do not have somatic mutations and copy number changes (*31*). Thus, building a translational dependency map using expression-based modeling robustly predicts gene essentiality and offers a flexible framework that can be broadly applied across malignant and normal tissue types alike.

**Figure 2.**
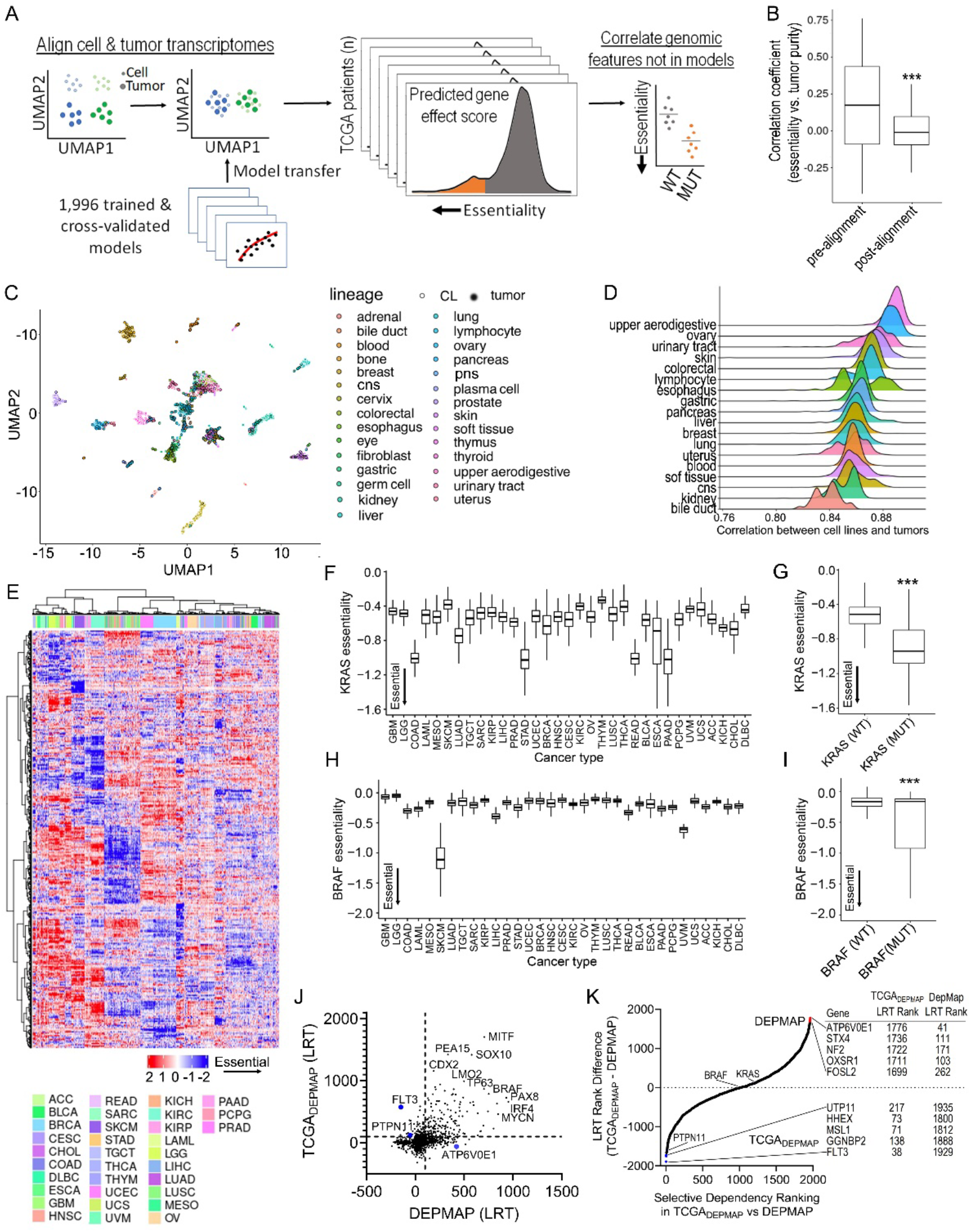
Building a translational dependency map: TCGADEPMAP. (**A**) Schematic of gene essentiality model transposition from DEPMAP to TCGA, following alignment of genomewide expression data to account for differences in homogeneous cultured cell lines and heterogenous tumor biopsies with stroma. (**B**) Correlation coefficients of gene essentiality scores and tumor purity before and after transcriptional alignment. (**C**) UMAP visualization of normalization of genomewide transcriptomes improves alignment between cultured cells and patient tumor biopsies with contaminating stroma. (**D**) Correlation coefficients of essentiality profiles of different lineages of cultured cell models and TCGA patient tumors. (**E**) Unsupervised clustering of predicted gene essentiality scores across TCGADEPMAP revealed strong lineage dependencies. Blue indicates gene genes with stronger essentiality, and red indicates genes with less essentiality. (**F**) *KRAS* dependency was enriched in TCGADEPMAP lineages with high frequency of *KRAS* gain-of-function (GOF) mutations, including colon adenocarcinoma (COAD), lung adenocarcinoma (LUAD), stomach adenocarcinoma (STAD), rectal adenocarcinoma (READ), esophageal carcinoma (ESCA), and pancreatic adenocarcinoma (PADD). (**G**) *KRAS* essentiality correlated with *KRAS* mutations in all TCGADEPMAP lineages. **(H)** *BRAF* dependency in TCGADEPMAP was enriched in skin cutaneous melanoma (SKCM), which has a high frequency of GOF mutations in *BRAF*. (**I**) *BRAF* essentiality correlated with *BRAF* mutations in all TCGADEPMAP lineages. (**J**) Scatter plot of model selectivity in TCGADEPMAP and DEPMAP, as determined by normality likelihood (NormLRT). (**K**) Ranking of model selectivity between in TCGADEPMAP and DEPMAP, as determined by the NormLRT scores. \*\*\**P* < 0.001, as determined by the Wilcoxon rank sum test for two group comparison and Kruskal-Wallis followed by Wilcoxon rank sum test with multiple test correction for the multi-group comparison.

As outlined in **Figure 2A**, the expression-based predictive models of DepMap dependencies were transposed to the transcriptomic profiles of 9,596 TCGA patients, following alignment to account for differences between the expression profiles of homogenous cell lines and tumor biopsies with varying stromal content. To control for the confounding stromal signatures, the expression profiles of the DEPMAP cell lines and TCGA patients were transformed by contrast PCA (cPCA) to remove cPC1-4 that comprised the tumor stroma (*34*) and correlated with tumor purity (**Figure 2B** and **Table S4**). The removal of cPC1-4 significantly improved the alignment of the expression-based dependency models when applied to TCGA patient transcriptomes (**Figure 2C, D** and **Figure S1**). Unsupervised clustering of gene essentialities across TCGA_DEPMAP_ revealed striking lineage dependencies (**Figure 2E** and **Table S5**), including well-known oncogenes such as *KRAS* (**Figure 2F, G**) and *BRAF* (**Figure 2H, I**). For example, *KRAS* essentiality was markedly stronger in *KRAS*-mutant STAD, READ, PAAD, and COAD lineages (**Figure 2F, G**), whereas *BRAF* essentiality was strongest in *BRAF*-mutant SKCM (**Figure 2H, I**). To ensure that the associations between dependencies and mutations were not due to the same underlying predictive features, the accuracy of elastic-net models to predict essentiality and somatic mutations in the same genes were compared. The comparison was restricted to genes with cross-validated models of essentiality and somatic mutations with >2% prevalence (n = 891 models). The elastic-net models were allowed to select the most informative predictive features for mutation and essentiality for each gene, as the best predictors for essentiality may not be the best features to predict mutation. Comparison of the AUCs of the two model sets revealed that transcriptomic features were significantly more predictive of gene essentiality compared with mutational status (**Figure S2**). Considering that the expression-only models of essentiality did not include genomic features, these data further demonstrate that the essentiality scores in TCGA_DEPMAP_ can be independently correlated with genomic features in patient tumors.

Strongly selective dependencies (SSDs) have also been characterized in cell-based maps using the normality likelihood ratio test (NormLRT) to rank whether an essentiality fits a normal or t-skewed distribution (i.e., selective) across the cohort (*20, 32*). A strength of this approach is the ability to rank SSDs regardless of the underlying mechanisms of dependency (e.g., lineage, genetic, expression, etc.). To compare the SSDs in cancer patients and cell models, NormLRT was applied to gene effect scores for the cross-validated essentiality models in TCGA_DEPMAP_ and DEPMAP, respectively. Most SSDs (NormLRT > 100) correlated well between TCGA_DEPMAP_ and DEPMAP (*r* = 0.56, p < 0.0001), including *KRAS, BRAF, MYCN*, and many other known SSDs (**Figure 2J** and **Table S6**). Although most SSDs correlated well between TCGA_DEPMAP_ and DEPMAP, there were several examples where the SSDs differed between patients and cell models (**Figure 2I**). Notably, the druggable oncogenes (e.g., *FLT3* and *PTPN11*) were more prominent SSDs in TCGA_DEPMAP_ patients than DEPMAP cell lines, whereas other notable SSDs in the DEPMAP (e.g., ATP6V0E1) were less noticeable in TCGA_DEPMAP_ (**Figure 2J, K**). The top predictive features for essentiality of *FLT3* (self-expression) and *ATPV6V0E1* (paralog expression) did not differ between DEPMAP and TCGA_DEPMAP_, yet the distribution and prevalence of strong dependency scores varied across lineages between patients and cell lines (**Figure S3A**-**D**). Likewise, the dependency on *PTPN11* (SHP2) was noticeably more selective in TCGA_DEPMAP_ than DEPMAP (**Figure 3J, K**), which was reflected by greater essentiality in a subset of breast cancer patients (**Figure S3E**) that was absent from breast cancer cell lines (**Figure S3F**). A Fisher’s exact test of the genetic drivers that were enriched in TCGA_DEPMAP_ breast cancer patients that were most dependent on *PTPN11* included *TP53* mutations and *HER2* amplifications (**Figure S3G**), whereas *FAT3* deletions and *GATA3* mutations were significantly depleted in these patients (**Figure S3H**). Particularly in the case of *ERRB2*, which signals through *PTPN11* and the RAS pathway, these data fit with the observation that RAS pathway inhibition, including SHP2 inhibitors, are more potent in the 3D versus 2D context (*33, 34*). Thus, the presence of TCGA_DEPMAP_ breast cancer patients that were highly dependent on *PTPN11* is likely due to the 3D context of patient tumors, whereas DEPMAP breast cancer cell lines with similar genetic drivers are not *PTPN11* dependent due to the 2D context of cultured cells. Collectively, these data demonstrate that identifying SSDs can be impacted by different prevalence and distributions of the underlying drivers in patients and cell models, which can be overcome by patient-relevant dependency maps, such as TCGA_DEPMAP_.

**Figure 3.**
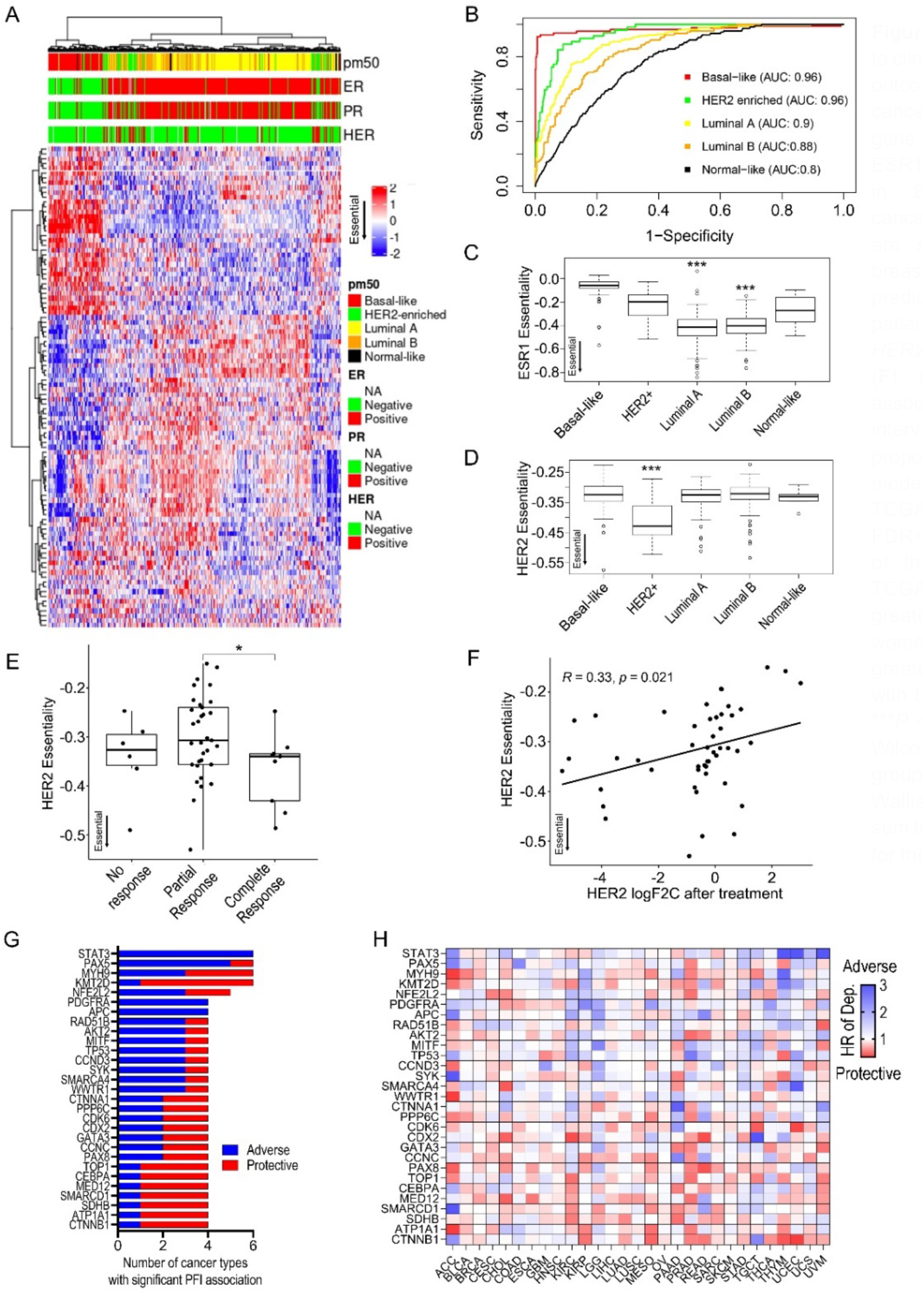
Translating TCGADEPMAP to clinically relevant phenotypes and outcomes. (**A, B**) Subtyping breast cancer dependencies by the top 100 gene dependencies (DEP100). (**C**) ESR1 dependencies are strongest in ER-positive luminal breast cancer. (**D**) *HER2* dependencies are strongest in *HER2*-amplified breast cancer. *HER2* dependency predicts trastuzumab response in patients (**E**) and correlates with *HER2* expression after treatment (**F**). (**G)** Top gene essentialities associated with the progression free interval (PFI) by univariate Cox-proportional hazard regression model across multiple lineages in TCGADEPMAP (Benjamini-Hochberg, FDR<0.2). (**H**) Hazard ratios (HR) of the top essentialities across TCGADEPMAP. Blue indicates a greater dependency associated with worse outcome and red indicates a greater dependency is associated with better outcome. \**P* < 0.05 and \*\*\**P* < 0.001, as determined by the Wilcoxon rank sum test for two group comparison and Kruskal-Wallis followed by Wilcoxon rank sum test with multiple test correction for the multi-group comparison.

### Translating TCGA_DEPMAP_ to clinically relevant phenotypes and outcomes

Another strength of translational tumor dependency maps is the ability to assess the impact of gene essentiality on clinically relevant phenotypes, such as molecular subtyping, therapeutic response, and patient outcomes. To evaluate the utility of TCGA_DEPMAP_ for therapy-relevant patient stratification, an unsupervised clustering of the 100 most variable gene dependencies was performed using the TCGA_DEPMAP_ breast cancer cohort (**Figure 3A**). The 100-dependency signature (DEP100) performed comparably to the established PAM50 signature (*35*) in classifying breast cancer subtypes (AUC close to 0.9 for most subtypes), despite only 3 overlapping genes between PAM50 and DEP100 (**Figure 2B**). Dependency subtyping with DEP100 predicted significantly higher *ESR1* essentiality in ER-positive tumors (**Figure 3C**) and higher *HER2* essentiality in *HER2*-amplified tumors (**Figure 3D**). Finally, due to the limited accessibility of therapeutic response data in TCGA (*36*), we applied the predictive gene essentiality model for *HER2* to a clinical trial of *HER2*-amplified breast cancer patients receiving trastuzumab (*37*), which revealed significantly higher predicted *HER2* dependency in patients that responded better to trastuzumab (**Figure 3E, F**). Taken together, these data establish the physiological relevance of TCGA_DEPMAP_ to associate dependencies with common clinicopathological features, such as molecular subtyping and therapeutic response.

The ability to associate gene essentiality with patient survival is a unique strength of TCGA_DEPMAP_, which is not accessible using cell-based dependency maps. Moreover, outcomes driven by perturbations of oncogenic pathways and genetic drivers of human cancers are likely not captured by gene expression alone and rather require a readout of gene essentiality. To test this possibility, the cross-validated gene essentiality models (n = 1,996) were tested for association with the progression free interval (PFI) in TCGA_DEPMAP_. Among 29 cancer lineages that are well powered for PFI analysis (*36*), 105 known genetic drivers of human cancer were significantly associated with the PFI of TCGA patients (**Table S7**), including 29 that were prognostic in at least four cancer lineages (**Figure 3G**). For example, a stronger dependency on the druggable oncogene, *STAT3* (*38*), was significantly associated with a shortened time to disease progression of six different cancers (**Figure 3H**). Likewise, multiple other prevalent genetic drivers of human malignancies were associated with a significantly shorter PFI, including *PAX5* and *PDGFRA* (**Figure 3H**). Notably, both proteins have been investigated previously as prognostic indicators of poor outcomes by expression analysis in patient biopsies (*39, 40*), yet to our knowledge this is the first time that dependency on these oncogenes has been associated with worse outcome in patients using a translational dependency map.

### Using TCGA_DEPMAP_ to translate synthetic lethalities in human cancer

In addition to illuminating lineage and oncogenic dependencies, the DEPMAP has dramatically expanded the list of potential synthetic lethalities (i.e., the loss of a gene sensitizes tumor cells to inhibition of a functionally redundant gene within the same pathway) (*6, 16, 17, 41, 42*). However, one of the current limitations of the DEPMAP is that the available cancer cell models do not yet fully recapitulate the genetic and molecular diversity of TCGA patients (*25*). Thus, we assessed the landscape of predicted synthetic lethalities with LOF events (damaging mutations or deletions) in TCGA_DEPMAP_. Lasso regression analysis of gene essentiality profiles and 25,026 LOF events detected in TCGA_DEPMAP_ yielded 633,232 synthetic lethal candidates (FDR<0.01), which were too numerous to experimentally validate by conventional methods. To prioritize the synthetic lethal candidates, the gene interaction scores were correlated with the mutual exclusivity of corresponding mutations in TCGA_DEPMAP_, which narrowed the list to 28,609 candidates (FDR<0.01). Multiple additional criteria were then applied to refine the list further by enriching for predicted paralogs with close phylogenic distance to prioritize candidates with redundant functions due to sequence homology. All told, this approach identified many known synthetic lethal pairs (e.g., *STAG1/2, SMARCA2/4*, and *EP300/CREBBP*) (*43*–*45*) and previously untested synthetic lethal candidates, demonstrating that TCGA_DEPMAP_ is well-powered to predict synthetic lethal relationships with LOF events in patient tumor biopsies (**Figure S4A**-**C** and **Table S8**).

Synthetic lethalities that were predicted with LOF events in the TCGA_DEPMAP_ (n = 604 pairs) were experimentally tested using a novel multiplexed CRISPR/AsCas12a screening approach across representative cell models of five cancer lineages (**Figure 5A, B**). Additional pairs (n = 261 controls) were added to the library to control for screen performance, including essential paralog pairs and non-essential pairs of TSGs and interacting partners (**Table S8**). An initial pilot screen was performed using five cancer cell models, which experimentally validated 69 TCGA_DEPMAP_ synthetic lethalities in at least one representative cell model (**Table S9**). As these data were being generated, an enhanced AsCas12a (enAsCas12a) enzyme was reported that is compatible with CRISPR/AsCas12a libraries (*46*), enabling replication of the initial pilot screens and expansion to a total sixteen total cancer cell models. Notably, the replication of the initial screens was highly concordant across the five cell models in common (average *r* = 0.69) (**Figure S5**), as well as detection of increased depletion of essential controls and synthetic lethal partners compared with non-essential controls (**Figure 4C**). In addition to novel pairs, multiple known synthetic lethalities (*HSP90AA1/B1* (*47*), *DDX19A/B* (*47*), *HDAC1/2* (*47, 48*), *SMARCA2/4* (*47, 48*), *EP300/CREBBP* (*45*), *STAG1/2* (*44, 48*) were replicated across multiple cell lines in both cohorts (**Table S9**), demonstrating the robustness of the multiplex CRISPR/Cas12a screening platform to test synthetic lethalities. For example, screening of H1299 cells confirmed seventeen synthetic lethalities that were predicted by TCGA_DEPMAP_, including multiple known pairs and several novel synthetic lethalities, such as *PAPSS1/2* (**Figure 4D**). Of the 604 synthetic lethalities predicted by TCGA_DEPMAP_, a total of 78 (13%) were experimentally validated in at least one representative cell model (**Figure 4E** and **Table S9**). Notably, as observed elsewhere (*41, 43, 48*), the sensitivity to synthetic lethalities varied between cell models and lineages, implicating the prevalence of unknown modifiers of synthetic lethality that manifest in different cellular contexts and are yet to be fully understood.

**Figure 4.**
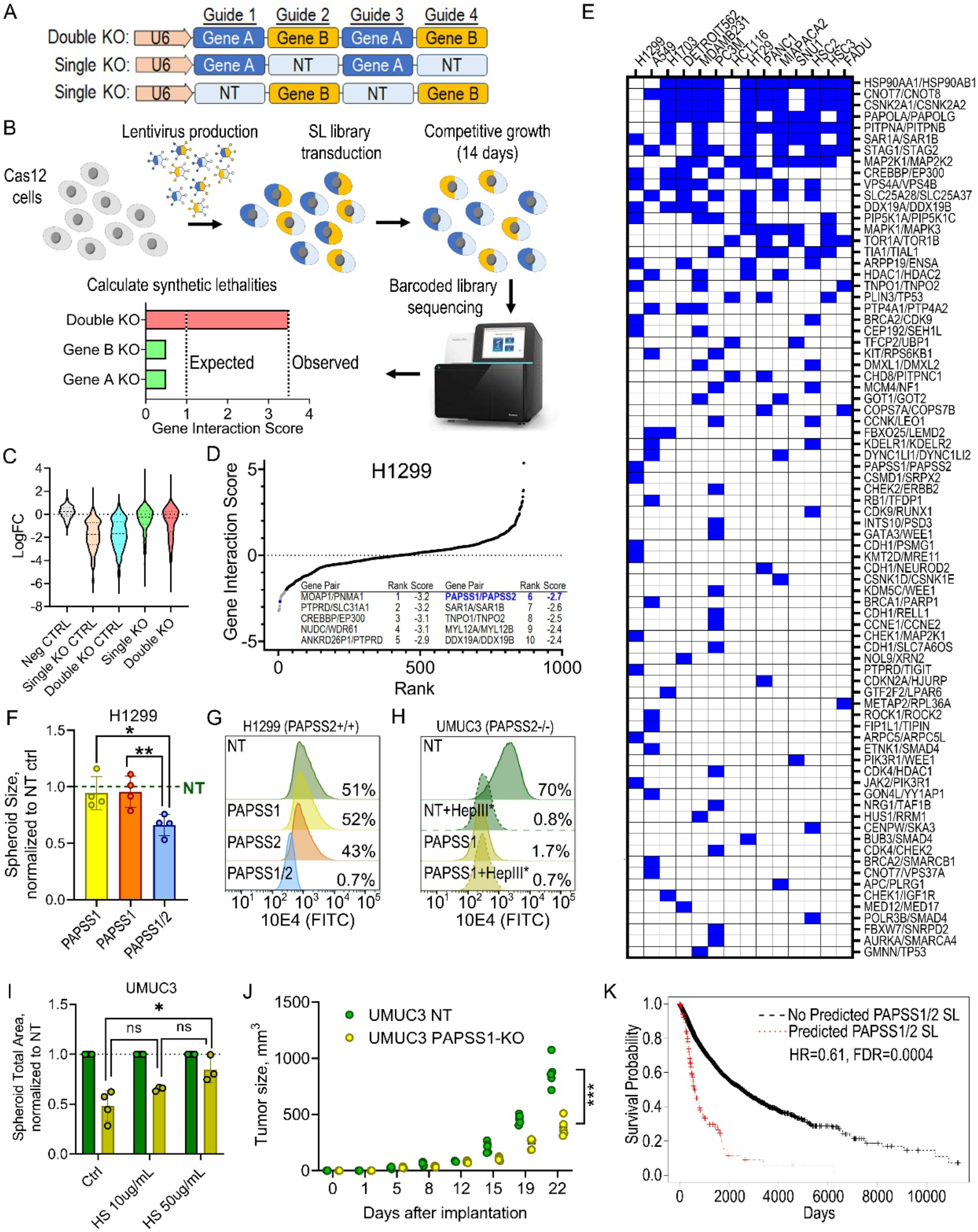
Using TCGADEPMAP to translate synthetic lethalities in human cancer. (**A**) Schematic of the CRISPR/Cas12 library multiplexed guide arrays targeting one or two genes per array. (**B**) Schematic of the synthetic lethality screening approach using the CRISPR/Cas12 library. (**C**) Violin plots of target-level CRISPR of log2 fold-changes for non-targeting guide (Neg CTRL), single knockout guides targeting essential genes (Single KO CTRL), double knockout guides targeting essential genes (Double KO CTRL), single knockout guides of TCGADEPMAP candidates (Single KO), and double knockout guides of TCGADEPMAP candidates (Double KO). (**D**) Rank plot of target level gene interaction (GI) scores in H1299 cells, including the top ten synthetic lethalities (table insert). The novel synthetic lethality, *PAPSS1/2*, is highlighted in blue. (**E**) Distribution of synthetic lethal candidates from TCGADEPMAP with experimental evidence of synthetic lethality in the CRISPR/Cas12 multiplexed screening across 14 cancer cell lines. A blue box indicates a GI score < -2. (**F**) Spheroid size of H1299 cells with single or dual *PAPSS1* / *PAPSS2* knockouts, normalized to non-targeting (NT) control spheroids. (**G**) Flow cytometry histogram overlay plots of viable H1299 cells (DAPI-) showing loss of cell surface sulfonated heparan sulfate proteoglycans (HSPGs) as detected by antibody 10E4-GFP in *PAPSS1/2*-DKO but not control. cells. Percentages shown are frequency of parent of positive expression. (**H**) Flow cytometry plots as in (**G**). HepIII* indicates that cells were enzymatically treated with Heparinase III, an enzyme that specifically cleaves sulfonated HS chains. (**I**) Bar diagram showing spheroid size of UMUC3 *PAPSS1*-KO (yellow) normalized to NT (green), either untreated (Ctrl) or supplemented with 10ug/ml and 50ug/ml of exogenous Heparan Sulfate (HS). (**J**) Diagram showing tumor sizes over time after *in vivo* implantation of 1e6 UMUC3 NT or *PAPSS1*-KO cells in SCID/Beige mice. Each dot is an individual mouse. (**K**) Kaplan-Meier plot of TCGADEPMAP patients with a predicted *PAPSS1/2* synthetic lethality have worse outcome compared with the rest of the cohort, as determined by a Cox log-rank test.

**Figure 5.**
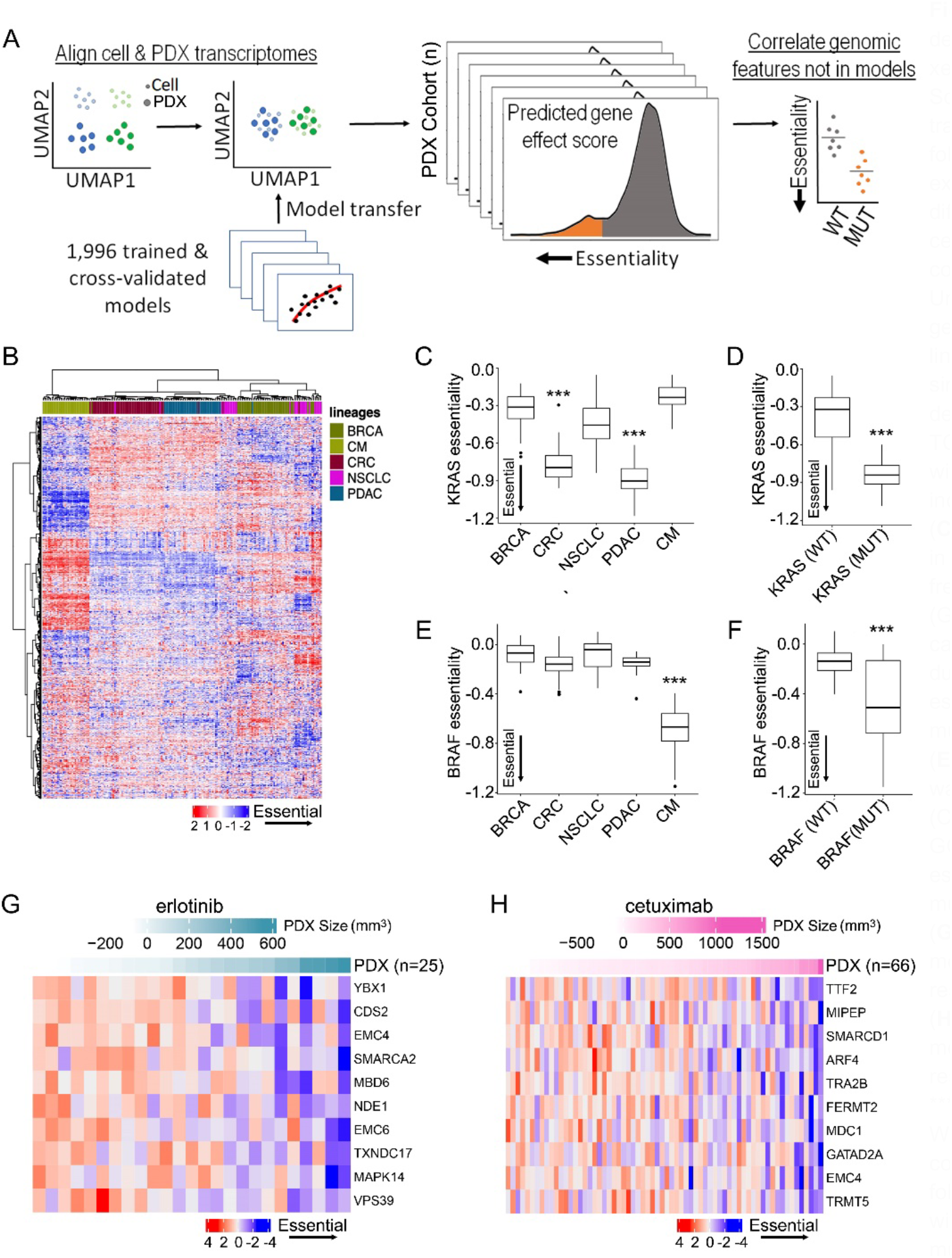
Building a translational dependency map in patient-derived xenografts: PDXEDEPMAP. **A**) Schematic of gene essentiality model transposition from DEPMAP to PDXE, following alignment of genomewide expression data to account for differences in homogeneous cultured cell lines and PDX samples with contaminating stroma. (**B**) Unsupervised clustering of predicted gene essentiality scores across five lineages in PDXEDEPMAP confirmed similar lineage drivers of gene dependencies, as observed in TCGADEPMAP. Blue indicates genes with stronger essentiality, and red indicates genes with less essentiality. (**C**) *KRAS* dependency was enriched in PDXEDEPMAP lineages with high frequency of *KRAS* gain-of-function (GOF) mutations, including colorectal carcinoma (CRC) and pancreatic ductal carcinoma (PDAC). (**D**) *KRAS* essentiality correlated with *KRAS* mutations in all PDXEDEPMAP lineages. (**E**) *BRAF* dependency in PDXEDEPMAP was enriched in cutaneous melanoma (CM), which has a high frequency of GOF mutations in *BRAF*. (**F**) *BRAF* essentiality correlated with *BRAF* mutations in all TCGADEPMAP lineages. Top correlated gene essentiality models that correlate with PDX response to erlotinib in PDXEDEPMAP. Top correlated gene essentiality models that correlate with PDX response to cetuximab in PDXEDEPMAP. \*\*\**P* < 0.001, as determined by the Wilcoxon rank sum test for two group comparison and Kruskal-Wallis followed by Wilcoxon rank sum test with multiple test correction for the multi-group comparison.

One novel discovery using TCGA_DEPMAP_ was the prediction of *PAPSS1* synthetic lethality with deletion of *PAPSS2* (**Figure S4E)**, and the neighboring tumor suppressor, *PTEN* (**Figure SGB**), which are frequently co-deleted in TCGA patient tumors (43% co-incidence) (**Figure S4H-J**). *PAPSS1/2* are functionally redundant enzymes essential for synthesis of 3’-phosphoadenosine 5’-phosphosulfate (PAPS) which is required for all sulfonation reactions (*49*), suggesting that loss of *PAPSS1/2* is synthetic lethal due to the inability to sulfonate proteins. To test this hypothesis, *PAPSS1/2* were targeted in H1299 spheroids by ribonucleoprotein (RNP), followed by measurement of spheroid growth and sulfonation levels of heparan sulfate proteoglycan (HSPG) chains on the cell surface by flow cytometry. Confirming the CRISPR/Cas12 screen data (**Figure 4D**), dual loss of PAPSS1/2 significantly reduced H1299 spheroid growth compared with controls (**Figure 4F** and **S6A**-**B**), which coincided with loss of HSPG sulfonation (**Figure 4G**). Likewise, targeting *PAPSS1* by RNP in UMUC3 cells, which endogenously lack *PAPSS2* and *PTEN*, also significantly depleted HSPG sulfonation and coincided with significant spheroid growth reduction (**Figure 4H** and **S6A**-**B**), which could be rescued by addition of exogenous heparan sulfate (**Figure 4I**). Finally, *PAPSS1/2* synthetic lethality was confirmed *in vivo*, as demonstrated by a significant tumor growth reduction of UMUC3 tumors without *PAPSS1/2* compared with control tumors lacking only *PAPSS2* (**Figure 4J** and **S6C**). Collectively, these data demonstrate that translational dependency maps, such as the TCGA_DEPMAP_ are powerful tools to uncover previously underrepresented synthetic interactions in cancer models that are likely to be patient relevant.

TCGA_DEPMAP_ is unique in its ability to uncover potential synthetic lethalities that can be related to patient outcomes, enabling the prioritization of the experimentally validated synthetic lethalities that correlate with the worst outcome and therefore likely to have the greatest clinical impact if druggable. To test this possibility, a Cox log-rank test was used to assess overall survival (OS) of TCGA patients that correlated with predicted gene essentiality by TCGA_DEPMAP_ and LOF events (mutation, deletion, or both) of the putative synthetic lethal partner. After controlling for tumor lineage, *PAPSS1* dependency in TCGA_DEPMAP_ was correlated with significantly worse OS (HR=0.63, p=0.0003) in patients with *PAPSS2* deletion (**Figure 4K**), demonstrating that *PAPSS1* is a novel synthetic lethality target with potentially high translational impact. Collectively, these data demonstrate for the first time that translational dependency maps can enable the discovery, validation, and translation of novel synthetic lethalities.

### Building a translational dependency map in patient-derived xenografts: PDXE_DEPMAP_

In addition to building TCGA_DEPMAP_, a similar approach was applied to generating an orthogonal translational dependency map generated using the PDX Encyclopedia (PDXE_DEPMAP_) (*29*). As outlined in **Figure 5A**, PDXE_DEPMAP_ was assembled by transferring the cross-validated 1,996 expression-only models from the DEPMAP to the PDXE (n = 191 tumors) using the aligned genomewide expression profiles from the PDXE (**Table S10**). Unsupervised clustering of gene essentialities across five well-represented lineages in PDXE_DEPMAP_ confirmed that lineage is a key driver of gene dependencies (**Figure 5B**), fitting with the observations made in TCGA_DEPMAP_ (**Figure 2E**). PDXE_DEPMAP_ also detected markedly stronger *KRAS* essentiality in *KRAS*-mutant PDX of PDAC and CRC lineages (**Figure 5C, D**), while *BRAF* essentiality was strongest in *BRAF*-mutant PDX of CM (**Figure 5E, F**). These data collectively demonstrate that the PDXE_DEPMAP_ performed comparably to TCGA_DEPMAP_ and is well powered to detect gene essentiality signals in PDX models.

In addition to orthogonal validation of TCGA_DEPMAP_, a unique strength of PDXE_DEPMAP_ is the ability to assess gene essentiality in the context therapeutic responses across five cancer lineages and 15 molecular therapies (*29*). To test the ability of gene essentiality to predict the response to corresponding targeted therapies, the change in PDX burden from baseline to experimental endpoint was correlated with target gene essentiality. This revealed that 80% of drugs (12 of 15) were significantly correlated (p<0.05) with the predicted essentiality of the target gene (**Table 1**). For example, trastuzumab response in the PDXE_DEPMAP_ was strongly predicted by *HER2* dependency (R=0.4849, p=0.002, AUC=0.75), in line with the predictive power of *HER2* dependency on trastuzumab responsiveness in patients with *HER2*-amplified breast cancer (**Figure 3E, F**). Other examples, such as erlotinib (R=0.4937, p=0.01, AUC=0.78) and cetuximab (R=0.2293, p=0.06, AUC=0.83), target the same gene (*EGFR*), providing the opportunity to explore dependency mechanisms of therapeutic resistance across modalities. Comparisons of PDX responses to erlotinib or cetuximab revealed dependencies within two common pathways: the SWI/SNF complex (*SMARCA2* and *SMARCAD1*) and protein trafficking (*EMC4, EMC6, VPS39*, and *MAPK14*) (**Figure 4G, H**). Notably, components of both pathways have been implicated in resistance to *EGFR* inhibitors (*50, 51*), suggesting that targeting these dependencies would likely improve patient outcomes. Taken together, these data demonstrate the ability of gene essentiality to predict therapeutic response and highlight the translatability of PDX modeling to patient-relevant clinical outcomes.

**Table 1.**
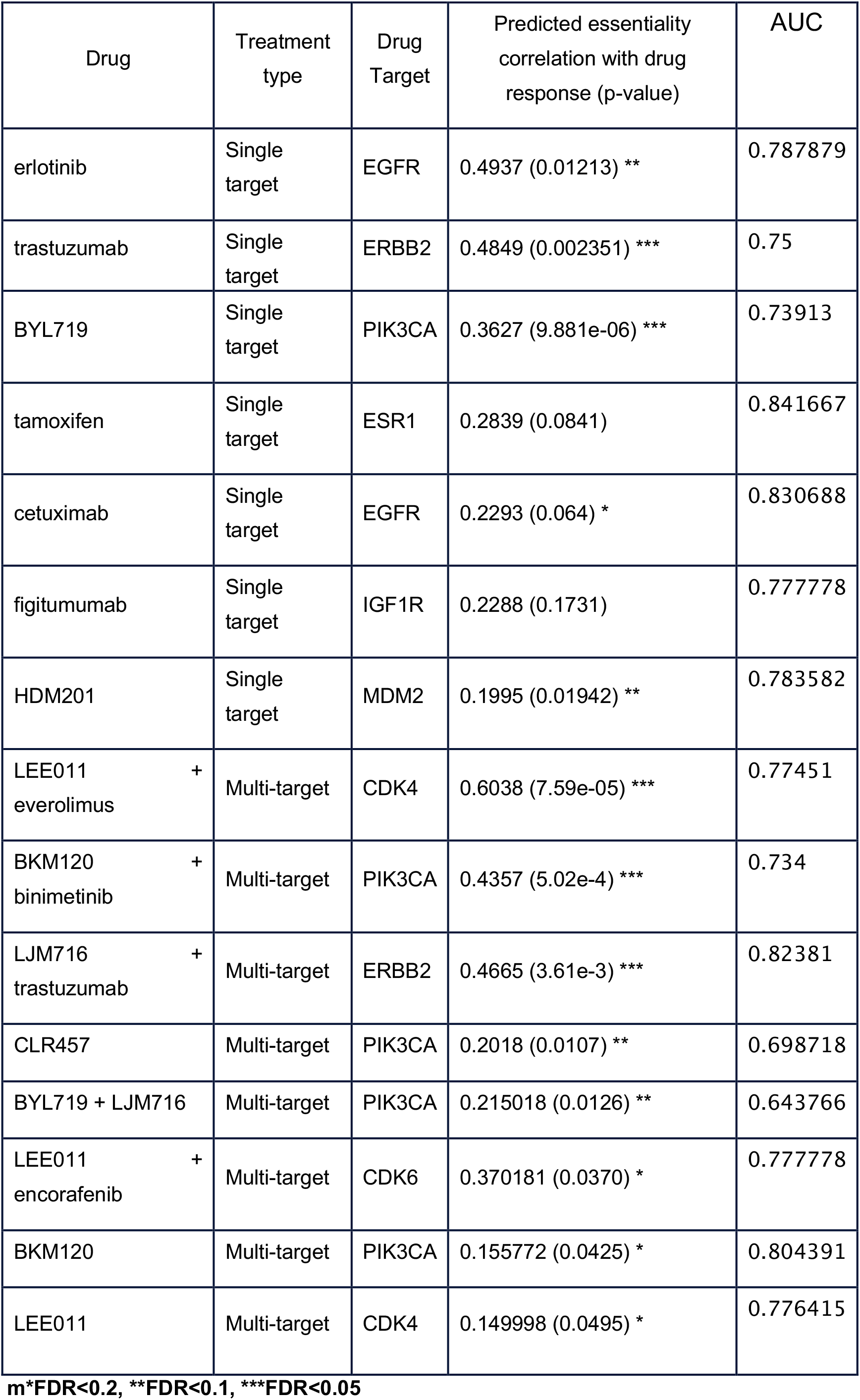
Correlation of gene essentiality and drug response in PDXE_DEPMAP_

### Building a translational dependency map in normal tissues: GTEX_DEPMAP_

A final objective of this study was to define gene essentiality in the context of normal tissues, which would provide a novel resource for prioritizing tumor dependencies with the best predicted tolerability. To achieve this objective, the expression-based dependency models from the DEPMAP were transposed using the aligned expression data from GTEX (GTEX_DEPMAP_), a compendium of deeply-phenotyped normal tissues collected from postmortem healthy donors (n=948) (*31*) (**Figure 6A** and **Table S11**). To assess the sensitivity of GTEX_DEPMAP_ to dependencies with low tolerability, the molecular targets of drugs with reported toxicities in the liver and blood (n=241) were compared across GTEX_DEPMAP_ (**Table S12**). This revealed that the average essentiality was significantly higher in liver and blood than other normal tissues. (**Figure 6B**). Likewise, unsupervised clustering of the 1,996 cross-validated gene essentiality models revealed strong tissue-of-origin dependencies in normal organs (**Figure 6B**), suggesting that tissue-specific biological context also contributes to gene essentiality in normal physiological settings. Taken together, these data demonstrate that GTEX_DEPMAP_ is sensitive to known toxicities, which cluster around different normal organ types.

**Figure 6.**
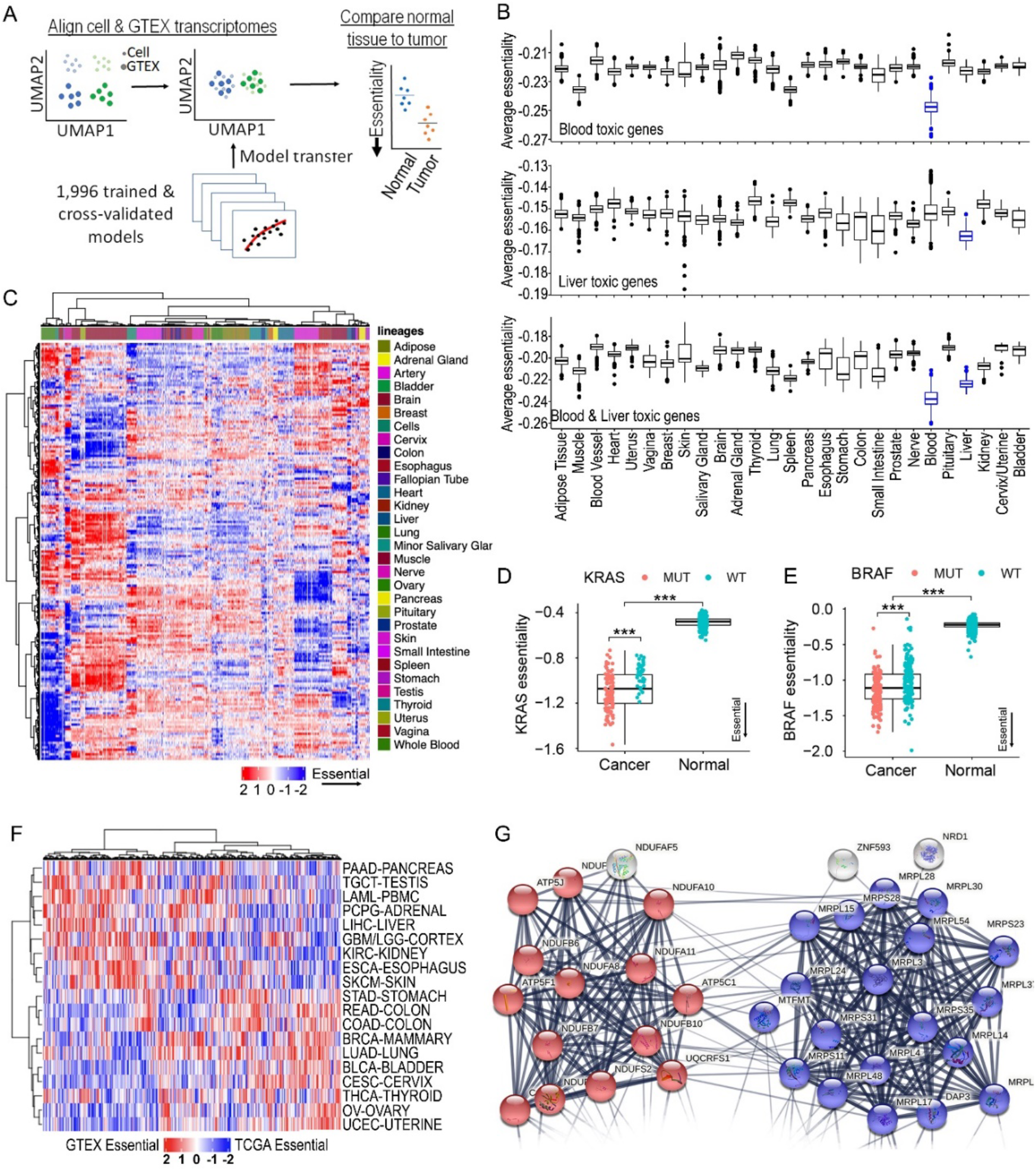
Building a translational dependency map in normal tissues: GTEXDEPMAP. (**A**) Schematic of gene essentiality model transposition from DEPMAP to GTEX, following alignment of genomewide expression data to account for differences in homogeneous cultured cell lines and normal tissue biopsies. (**B**) Average gene essentiality profile across normal tissues of GTEXDEPMAP for molecular targets with known liver and blood toxicities (in blue). (**C**) Unsupervised clustering of predicted gene essentiality scores across normal tissues. Blue indicates genes with stronger essentiality, and red indicates genes with less essentiality. (**D**) *KRAS* essentiality is significantly higher in PAAD with GOF mutations compared with normal pancreas in GTEXDEPMAP. (**E**) BRAF essentiality is significantly higher in SKCM with GOF mutations compared with normal skin GTEXDEPMAP. (**F**) Global differences between the predicted target efficacy score (TCGADEPMAP) and the normal tissue-of-origin tolerability score (GTEXDEPMAP). (**G**) STRING network analysis of the top 100 LUAD targets with the greatest predicted tolerability in normal lung reveals significant connectivity (p<1×10^−16^) and gene ontology enrichment oxidative phosphorylation (blue colored spheres; p=5.8×10^−11^) and mitochondrial translation (red colored spheres; p=2.9×10^−20^). \*\*\**P* < 0.001, as determined by the Wilcoxon rank sum test for two group comparison and Kruskal-Wallis followed by Wilcoxon rank sum test with multiple test correction for the multi-group comparison.

Comparing essentiality scores of known druggable oncogenes in TCGA_DEPMAP_ with GTEX_DEPMAP_ revealed greater dependency in malignant tissues versus the normal tissue-of-origin. For example, *KRAS* and *BRAF* essentialities appear to be concomitantly dependent on lineage and genetic drivers, as the normal tissues-of-origin were predicted to be significantly less affected in the GTEX_DEPMAP_ compared with TCGA_DEPMAP_ (**Figure 6D, E**). Likewise, similar observations were made for other oncogenic drivers that are approved therapeutic targets in cancer patients, such as *HER2* amplified breast cancer (**Figure S7**). In contrast, there was markedly less separation in the predicted essentialities of malignant tumors and normal tissues-of-origin for molecular therapies that have yet to be successful in clinical trials (**Table S13**). To refine the list of oncogenic pathways with significant differences in tumor efficacy and normal tissue-of-origin tolerability, we compared dependency (TCGA_DEPMAP_) and tolerability (GTEX_DEPMAP_) scores across all genes and tissues (**Figure 6F**). Pathway analysis of the strongest tumor dependencies with the least tissue-of-origin toxicity revealed enrichment of multiple oncogenic pathways and pathophysiological processes (**Table S14**, including dysregulation of oxidative phosphorylation (p=5.8×10^−11^) and mitochondrial translation (p=2.9×10^−20^) pathways that were enriched in LUAD compared with normal lung (**Figure 6G** and **S8**). Combined, these observations suggest that predicted gene essentiality in the context of a driver mutation and correspondingly low essentiality within the normal tissue-of-origin is likely to identify efficacious drug targets with acceptable tolerability.

## DISCUSSION

This study addresses the significant challenge of translating gene essentiality from cell-based dependency maps to the rapidly expanding catalogue of patient-relevant datasets. The translational dependency maps (TCGA_DEPMAP,_ PDXE_DEPMAP_, and GTEX_DEPMAP_) were built using expression-based predictive models of gene essentiality, because expression-only models performed comparably to multi-omics models that included genomic features (e.g., somatic mutations and copy number), as was reported elsewhere (*26, 27*). Another strength of expression-based predictive modeling is that it can be applied to the transcriptomic profiles of normal tissues (e.g., GTEX_DEPMAP_) that do not have appreciable levels of the somatic alterations that are observed in malignant tissues (*31*). Moreover, transcriptomics is frequently the most widely captured dataset for many largescale clinical studies (*52*), suggesting that the methods used here will be widely applicably across many studies. Indeed, the same expression-based predictive models recapitulated a similar dependency profile in TCGA_DEPMAP_ and PDXE_DEPMAP_ across different cancer lineages and genetic drivers. Both maps also accurately predicted therapy-relevant molecular subtypes, therapeutic responses, and disease outcomes, whereas GTEX_DEPMAP_ provided novel insights to normal tissue-of-origin tolerability. Collectively, these translational dependency maps offer clinically relevant aspects to gene essentiality that are not currently accessible in the traditional cell-based dependency maps.

### Translating tumor dependencies using TCGA_DEPMAP_ and PDXE_DEPMAP_

A challenge of cell-based dependency maps (e.g., DEPMAP) is the inability to fully recapitulate patient genomics, therapeutic responses, and many aspects of disease outcomes and patient survival (*36*). Using TCGA_DEPMAP_ to survey the landscape of translational dependencies in cancer patients, we observed a strong correlation between patient tumor type and corresponding cancer cell line lineages (**Figure 2C-E**). A comparison of predictive dependency models across TCGA_DEPMAP_ (**Figure 2E**) and PDXE_DEPMAP_ (**Figure 5B**) revealed that dependencies accurately predict tumor lineages (AUC=0.75), fitting with observations made in TCGA that tissue-of-origin dominates the molecular landscape of cancer (*53*). In addition to lineage dependencies, TCGA_DEPMAP_ and PDXE_DEPMAP_ also demonstrated the abilities of predicted dependencies to define therapy relevant molecular subtypes, including the ER-positive luminal breast cancer (**Figure 3A-C**) and *HER2*-amplified breast cancer (**Figure 3A-B, D**). Fitting with these observations, the response to trastuzumab and post-treatment *HER2* expression in patients with *HER2*-amplified breast cancer was significantly associated with predicted *HER2* essentiality (**Figure 3E**, **F**), as was the ability to predict trastuzumab response in PDXE_DEPMAP_ (**Table 1**). In total, PDXE_DEPMAP_ showed that the essentiality of a molecular target could be associated with the molecular therapy in 12 out of 15 cases (**Table 1**), as well as the ability to correlate dependencies with intrinsic resistance to therapeutic responses (**Figure 5G**, **H**). However, despite the novel findings, power analyses using these data also suggest that increased cohort sizes will be required to replicate and expand these findings. Thus, translational dependency maps combined with the expanding catalogue of patient genomics and PDX modeling (*3, 28*) are expected to provide novel molecular targets for patient-relevant therapies.

In addition to cancer lineage and subtype dependencies, TCGA_DEPMAP_ provided novel molecular insight to the genetic drivers of gene essentiality in patient populations. Notably, well-defined oncogenic drivers of tumor dependencies (e.g., *KRAS* and *BRAF*) were highly predictable in TCGA_DEPMAP_ and PDXE_DEPMAP_ (**Figure 2F-I** and **5C-F**), as would be expected based on the predictive response rates for molecular therapeutics targeting oncogenic drivers in patients (*54*). However, despite most SSDs correlating between TCGA_DEPMAP_ and DEPMAP, several notable dependencies showed different selectivity profiles between patients and cell models, including *FLT3, ATPV6V0E1*, and *PTPN11*. Some of these discrepancies appear to be attributed to cohort-specific distributions of the underlying drivers of SSDs (e.g., *FLT3* and *ATPV6V0E1*) (**Figure S3A**-**D**), whereas others were likely attributable to different pathophysiological contexts, such as the 3D contexts of intact tumors versus the 2D contexts of cultured cells (e.g., *PTPN11*) (**Figure S3E**-**H**). Importantly, TCGA_DEPMAP_ and PDXE_DEPMAP_ also enabled the association of tumor dependencies with clinical outcomes, including predicting therapeutic responses (**Figures 3E-F**, **5G-H**, and **Table 1**) and patient survival (**Figure 3G-H**). Taken together, these data demonstrate the value of translational dependency maps to interpret gene essentiality in patient-relevant contexts and extending these findings to predicting novel tumor dependencies that impact patient outcomes.

Multiple known synthetic lethalities were also detected in TCGA_DEPMAP_ (*STAG1/2, SMARCA2/4*, and *EP300/CREBBP*) (*44, 45, 47, 48*), as well as previously unappreciated candidate pairs that have not been reported elsewhere (e.g., *PAPSS1/2*). Intriguingly, *PAPSS2* deletion is prevalent in the TCGA, yet deletions of this genetic loci were mostly absent from DEPMAP cell lines. We hypothesize that loss of *PAPSS2* is likely driven by its proximity to PTEN and is an example of collateral deletion in patient tumors (*55*). Since *PTEN* is largely silenced by LOF mutations in cancer cell lines and not deletion, a synthetic lethality between *PAPSS1* and endogenous collateral deletion of *PAPSS2* with *PTEN* is likely undetectable in cell-based dependency maps. Nonetheless, increased dependency on *PAPSS1* was observed in one of the few DEPMAP cell models with endogenous *PAPSS2* and *PTEN* collateral deletion, UMUC3 (**Figure 4I, J**), which could be attributed to the inability of these cells to sulfonate proteins (**Figure 4H**). Importantly, the unique ability of TCGA_DEPMAP_ to associate synthetic lethal mechanisms with patient outcomes revealed a worse overall survival of patients with an endogenous loss of *PAPSS2* and a predicted synthetic lethality with *PAPSS1* dependency (**Figure 4K**). Thus, these data collectively highlight the benefits of translational dependency maps that closely match the pathophysiological contexts of intact patient tumors and the diversity of patient genomic datasets to identify clinically-relevant mechanisms (*1, 28*).

### Translating tolerability of dependencies using TCGA_DEPMAP_ and GTEX_DEPMAP_

As the landscape of gene essentiality continues to expand, a better understanding of the impact on normal tissues to predict tolerability is needed. To address this challenge, we applied the expression-based dependency models to GTEX, a compendium of deeply-phenotyped normal tissues collected from postmortem healthy donors (*31*). This novel map of gene essentiality in the context of normal tissues (i.e., GTEX_DEPMAP_) was able to predict greater gene essentiality in normal blood and liver for the molecular targets of multiple drugs with observed toxicities in those organs (**Figure 6B**). Thus, these data support that GTEX_DEPMAP_ is also well-powered to detect other genes with potential liabilities in other organs. Strikingly, unsupervised clustering of the predicted gene essentialities across 17,382 normal tissues in the GTEX_DEPMAP_ also demonstrated a significant impact of tissue-of-origin on gene essentiality (**Figure 6C**). One potential reason is that pathways required for maintaining normal tissue homeostasis are frequently co-opted for oncogenesis (*56*) and several such pathways serve as key therapeutic targets for cancer (*57*). In addition to lineage pathways, most successful cancer therapeutics are also targeted towards somatic drivers, which are absent from normal tissues and therefore likely to be more tolerable (*56, 57*). A unique aspect of this study was the ability to systematically compare gene essentiality associated with somatic mutations in TCGA_DEPMAP_ with the normal tissue-of-origin tolerability profiles in GTEX_DEPMAP_. Strong oncogenic dependencies with acceptable tolerability (e.g., *KRAS* and *BRAF*) (*56, 57*) revealed marked differences between predicted gene essentiality in malignant and normal tissues, and this therapeutic window was further widened in the presence of the oncogenic mutation (**Figure 6D-E**). Systematically expanding this analysis across all gene essentiality models in TCGA_DEPMAP_ and GTEX_DEPMAP_ revealed wide variability in the predicted tolerability windows, implicating the existence of other dependencies with strong genetic drivers that are likely to be more tolerable as therapeutic targets. However, when interpreting these data we also recommend exercising caution, as the tolerability windows predicted by comparing tissue-of-origin gene essentiality between TCGA_DEPMAP_ and GTEX_DEPMAP_ likely does not yet fully capture the other dose-limiting toxicities that pose challenges to clinical drug development (*58*). As such, future efforts to model gene essentiality in normal tissues should expand to incorporate systems approaches to integrating tolerability signals across multi-organ physiological pathways and systems.

### Comparison of TCGA_DEPMAP_ with other translational dependency maps

During the completion of this study, Chiu et al (*27*) took a complimentary approach to building a translational dependency map (DeepDEP) using deep learning (DL) from the integrative genomic, epigenomic, and transcriptomic profiles of TCGA patients and DEPMAP cell lines. Here, we used elastic-net regularized regression models of expression data for predicting gene essentiality and tolerability, as these expression-based models performed comparably to multi-omics models and can be applied to malignant tissue (TCGA_DEPMAP_ and PDXE_DEPMAP_) and nonmalignant tissues (GTEX_DEPMAP_). The DeepDEP authors also highlighted that a simplified DL model using expression only (Exp-DeepDEP) performed comparably well to DeepDEP (*27*), suggesting that both approaches are dominated by expression data (*27*). For lack of other ground truths, we compared the predicted tumor dependencies of TCGA_DEPMAP_ and DeepDEP by pan-cancer lineage and breast cancer subtypes, as these were annotated by TCGA and DEPMAP. Compared with DeepDEP, the predicted dependencies by TCGA_DEPMAP_ were more accurate in identifying cancer lineages and breast cancer subtypes (**Figure S9A**-**B**). Moreover, TCGA_DEPMAP_ was able to accurately cluster tumors and cell models of the same cancer lineage, whereas DeepDEP was comparably less able cluster tumors and cell lines by t-SNE, despite the dominance of expression in the DeepDEP dependency models (*27*). One potential explanation is the liability of DL models to overfit highly dimensional data (e.g., transcriptomics) (*59*), which was potentially compounded by the unlabeled pretraining stage of DeepDEP with TCGA expression data that included expression profiles of nonmalignant stroma (*27*). Indeed, adopting the approach by Warren et al (*60*) to remove the confounding effects of stromal signatures and tumor purity in patient biopsies resulted in more accurate transposition of the dependency models. Thus, the collective data demonstrated that the elastic-net models underlying TCGA_DEPMAP_, PDXE_DEPMAP_, and GTEX_DEPMAP_ performed well compared to DeepDEP, which is to our knowledge the only other translational dependency map to date. As additional studies become available, a more in-depth benchmarking of approaches for translating dependencies is warranted, including the ability to detect genetic drivers, synthetic lethalities, and other patient-relevant features.

### Future considerations for translational dependency maps

The translational dependency maps presented in this study (TCGA_DEPMAP_, PDXE_DEPMAP_, and GTEX_DEPMAP_) provide novel insights to gene essentiality and tolerability in the clinical context of patient tumors and normal tissues. The ability of these maps to accurately translate dependencies to patients is reliant on the ability to build predictive models from cell-based mapping, which is still at the early stages (∼1,000 mapped cell lines) and is expected to require 20X more data (∼20,000 mapped cell lines) to fully predict gene essentiality (*7*). Similar needs exist to expand and incorporate translational dependency maps to include compound screening efforts across large number of cell models, such as the PRISM (https://depmap.org/portal/) and GDSC initiatives (https://www.cancerrxgene.org/). Further, the observations that cell-based dependencies vary between 2D and 3D settings (*61*) and are impacted by crosstalk with the TME (*62*), suggests that gene essentiality is contextual and requires models with greater relevance to intact tumors, such as organoids. Likewise, it is equally plausible that accurately interpreting translational dependencies will require a deeper understanding of clonal heterogeneity with patient tumors that is lacking from homogenous cancer cell lines. To reach the full potential of translational dependency mapping, the catalog of patient genomic datasets will also likely require expansion to capture various stages of disease progression, including tumorigenesis (*2*), metastasis (*3, 63*), and therapeutic resistance (*3, 4, 63*). Furthermore, as precision cancer clinical trials continue to expand (e.g., MSK-IMPACT) (*4*), it will be increasingly possible to refine translational dependency maps by testing outcomes of molecular therapeutics with predicted target essentiality. The utility of translational “tolerability” maps in normal tissues (e.g., GTEX_DEPMAP_) remains to be fully explored and will likely benefit from further refinements to better capture aspects of dose-limiting toxicities (DLT) that impact drug development. To this end, we postulate that modeling gene tolerability could be best assessed in normal cell types by pairing CRISPR perturbations with single-cell RNA sequencing (*64, 65*) to broadly capture the alterations of pathways required for normal tissue homeostasis. Ultimately, we postulate that predictive modeling of dependency and tolerability in patients will increase the success of drug discovery by preemptively prioritizing targets with the best therapeutic index (i.e., high dependency and tolerability).

## MATERIALS AND METHODS

### Predictive modelling of gene essentiality using DEPMAP data

Two sets of elastic-net regression models were generated to predict gene essentiality from the DEPMAP (n = 897 cell lines) with RNA alone (i.e., expression-only) or combined with mutation and copy number profiles (i.e., multi-omics). Gene effect scores were estimated by CERES (*24*), which measures the dependency probability of each gene relative to the distribution of effect sizes for common essential and nonessential genes within each cell line (*25*). Because many genes do not impact cell viability (CERES<-0.5), elastic-net models were attempted only for genes with at least five dependent and non-dependent cell lines, which included 7,260 out of 18,119 genes (40%) with effects scores in the DEPMAP (1Q21 release). Genomewide datasets (19,005 genes) for RNAseq, mutations, and copy number variants (log2 relative to ploidy + 1) for the 897 cell lines were downloaded directly from the DEPMAP (1Q21, https://depmap.org/portal/). The *‘glmnet’* package (version 4.1.3) (*23*) was used to build elastic-net regularized regression models with balanced weights for L1 and L2 norm regularization. The alpha values were kept constant at 0.5 for all models. Models were 10-fold cross-validated using “lambda.min” from cv.glmnet from the glmnet R package (100 lambdas tested per model by default) to select the lambda showing the minimum error balanced with the prediction performance and the number of features selected, as described previously (*66*). The performance of the optimal model per was then assessed by Pearson’s correlation coefficient (R), with a “pass” threshold of R>0.2 and FDR<0.001 to correct for multiple hypothesis testing. Cross-validation confirmed 1,996 expression-only models and 2,045 multi-omics models, of which the majority of cross-validated models overlapped (n=1,797) between the two datasets (**Table S3**).

### Model transposition following transcriptional alignment of DEPMAP to TCGA, PDXE, and GTEX datasets to build TCGA_DEPMAP_, PDXE_DEPMAP_, and GTEX_DEPMAP_

The translational dependency maps TCGA_DEPMAP_, PDXE_DEPMAP_, and GTEX_DEPMAP_ were built using expression-only models of gene essentiality, based on relatively marginal performance gains in the multi-omics models of gene essentiality, as reported elsewhere (*26, 27*). To enable transposition of the cross-validated expression-only models (n=1,996) from the DEPMAP to TCGA (N = 9,596 tumors), PDXE (N=191 tumors), and GTEX (N=17,382 tissues across 54 tissues and 948 donors), the genomewide gene expression data sets were downloaded for TCGA (https://xenabrowser.net/datapages/), PDXE (*29*), and GTEX (https://gtexportal.org/home/datasets). For TCGA data, if multiple samples were collected from the same patient, only the primary tumor biopsy was included in TCGA_DEPMAP_. For GTEX, the potential biases introduced by sampling multiple organ tissues from each individual was assessed by UMAP analysis of the gene expression profiles across GTEX samples, which revealed that GTEX samples are clustered by tissue types rather than by individuals. Likewise, no evidence of clustering was observed based on other patient-specific clinical variables (e.g., cause of death, age, etc.), suggesting that the tissue-specific effects are the predominant drivers of gene expression in normal tissues.

Unsupervised cluster analyses by UMAP dimension reduction were used to evaluate the similarities in expression profiles of the DEPMAP cell lines compared with the tissue biopsies from TCGA, PDXE, and GTEX. As reportedly previously (*60*), the DEPMAP and TCGA expression profiles do not cluster well by UMAP alignment due to contaminating transcriptional profiles of stromal and immune cells, which would impact expression-based predictive modeling of gene essentiality. Likewise, UMAP clustering of expression profiles from DEPMAP cell line data compared with PDXE and GTEX samples revealed that transcriptional alignment of these data were equally problematic. To overcome this issue, expression data from DEPMAP and TCGA were quantile normalized and transformed by contrastive principal component analysis (cPCA) to identify misaligned components derived from the stromal contamination in TCGA. The top contrastive principal components (cPC1-4) were removed, followed by multiple-batch correction to normalize the expression data by matching the corresponding clusters in TCGA and DEPMAP. To assess transcriptional alignment on model transposition, the pre- and post-aligned TCGA_DEPMAP_ models were compared with tumor purity, which revealed a strong correlation between gene essentiality and tumor purity that was removed by transcriptional alignment (**Figure 2B**). An identical approach was utilized for aligning PDXE expression data, with the slight modification that only cPC1-3 required removal, as PDX models grown in immunocompromised mice lack the adaptive immune system and typically have lower stromal contamination. For aligning DEPMAP and GTEX data, a slightly different approach was used to combine quantile normalization and ComBat (*67*) to remove potential batch effects without using cPCA, as GTEX data only includes nonmalignant tissue.

### Characterization of TCGA_DEPMAP_

The distribution of the cross-validated expression-only models of gene essentiality (n=1,996) across lineages was assessed by unsupervised cluster analysis (Ward.D2 method) and visualized using the ComplexHeatmap R package (version 2.6.2). A similar approach was used for unsupervised cluster analysis and heatmap visualization for molecular subtyping of the breast cancer (BRCA) cohort of TCGA_DEPMAP_ using the top 100 most variable dependencies (DEP100) across BRCA cohort only. For lack of other ground truths, the performance of TCGA_DEPMAP_ to classify molecular subtypes of BRCA was benchmarked using a linear discriminant analysis (LDA) with leave-one-out cross-validation performed using the MASS package (version 7.3.51.4) for R and the CV=TRUE option in the function. Predictions for each cancer type and subtype was evaluated separately and the AUC values were determined using the function “roc” from the pROC (version 1.18.0) package for R and compared with the molecular typing and subtyping reported by the TCGA (https://www.cbioportal.org/) (*68*). In addition to BRCA molecular subtypes, a distinct subset of the 100 most variable dependencies from the pan-cancer TCGA_DEPMAP_ dataset was used to benchmark TCGA_DEPMAP_ more broadly, using an identical LDA with leave-one-out cross-validation, as described above. Finally, both analyses were repeated with the DeepDEP gene essentiality values reported by Chiu et al (*27*) and the receiver operating characteristic (ROC)–area under the curve (AUC) values were compared between TCGA_DEPMAP_ and DeepDEP predictions of cancer lineages and BRCA cancer subtypes (**Figure S9**).

Associations of dependencies with genomic features (somatic mutations and copy number variants) in TCGA_DEPMAP_ were assess using a Wilcoxon Rank Sum differential test as implemented using stat_compare_means function of ggpubr R package (version 0.4.0). The ability of expression features to predict essentiality and mutational status of same gene by elastic-net modeling was compared using the glmnet R package (version 4.1) with the same parameters for both model sets. The elastic-net models were allowed to select the most informative predictive features for mutation and essentiality for each gene, as the best predictors for essentiality may not be the best features to predict mutation. For AUC evaluation, we used -0.5 as cutoff for gene essentiality scores to determine sensitive and resistant cells for gene models. The AUC values are calculated using pROC R package (version 1.16.2). To characterize strongly selectivity dependencies (SSD), a normality likelihood ratio test (NormLRT) (*32*) was performed with slight modifications to rescale the larger NormLRT values observed in TCGA_DEPMAP_ due to a 10-fold larger cohort size (n=9,596) compared with DEPMAP (n=897). A bootstrapping of the DEPMAP gene effect scores was performed to estimate how the NormLRT scores change when scaling up from the DEPMAP cohort size (n=897 cell models) to the cohort size of TCGA (9,596). A linear fitting was performed to estimate the slope between DEPMAP and bootstrapped equivalent, which was as a scaling factor (0.07) to rescale TCGA NormLRT scores. Notably, outliers were identified based on the ranking NormLRT scores within each cohort, which therefore was not affected by the rescaling TCGA NormLRT scores. For TCGA breast cancer patients (n=765), we divided the patients into *PTPN11* dependent and non-dependent groups. The *PTPN11* dependent patients (77 patients) are selected as top 10% breast patients with lowest *PTPN11* essentiality scores. Among all the variants, we applied Fisher’s exact test for mutations with more than 5% frequency (12 mutations), deletions with more than 10% frequency (4,891 deletions) and amplifications with more than 10% frequency (4,831 amplifications). The test was performed using the fisher.test function in stats (version 4.0.3) R package with options ‘alternative=greater’ to calculate p-values for enrichment of variants for *PTPN11* dependent and non-dependent groups. The gene models (890 models) used for mutation predictions are selected from 1,996 cross-validated expression only essentiality model with mutation frequency over 2%.

### Associating clinical outcomes with tumor dependencies in TCGA_DEPMAP_

Due to the limited accessibility of therapeutic response data in TCGA (*36*), the association of *HER2* essentiality with response to trastuzumab (anti-HER2 antibody) was tested in a recent trastuzumab clinical trial of 50 *HER2*+ breast cancer patients with pre- and post-treatment biopsies that were analyzed by microarray (*37*), The microarray expression data were downloaded from NCBI GEO (accession # GSE76360) and the patient responses were defined by the study authors (*37*). Differences in predicted *HER2* essentiality in patients with different clinical responses was tested using ggpubr R package (version 0.4.0), followed by a Wilcoxon Rank Sum Test using stat_compare_means function in the package. Correlation of *HER2* essentiality and HER2 expression post treatment was tested by Pearson’s correlation, as calculated by stat_cor function ggpubr R package (version 0.4.0). Additionally, the correlation of TCGA_DEPMAP_ dependencies with the progression free interval (PFI) of TCGA patients was performed, excluding the LAML, DLBC, KICH, and PCPG cohorts based on the recommendations of Liu et al (*36*). The PFI data were directly downloaded from Liu et al (*36*) and the maximally selected rank statistics from the ‘maxstat’ R package was used to determine the optimal cut-point for dichotomization (high vs. low) of dependency scores (n=1,996 cross-validated models). The prognostic value of the resulting dichotomized dependency scores was evaluated using the log-rank test with FDR correction (Benjamini-Hochberg adjusted) to account for multiple hypothesis testing. The data were visualized by Kaplan–Meier curves and are interpreted as a hazard ratio (HR) > 1 indicated a worse expected outcome in patients with a higher dependency score at an FDR < 0.2.

### Predicting synthetic lethality relationships in TCGA_DEPMAP_

Multiple approaches were integrated to predict and prioritize synthetic lethality relationships with loss-of-function (LOF) events (defined as a predicted copy number loss or damaging mutation) in TCGA_DEPMAP_. Lasso regression was used to identify gene essentialities (n = 7,260 expression-only models) with increased dependencies associated with 25,026 LOF events in TCGA, as annotated by Bailey et al (*69*). For each model, the lambda value was selected as the lowest error by 5-fold cross-validation and the resulting models with coefficients >0.3 were further evaluated by t-test. The lasso regression analysis identified 633,232 predicted synthetic lethal candidates (FDR<0.01), which were too numerous to experimentally validate and required further prioritization. First, UNCOVER (*70*) was used to prioritize synthetic lethal candidates predicted by TCGA_DEPMAP_ that correlated with endogenous mutual exclusivity of LOF mutations (3% to 70% prevalence) in TCGA, with the hypothesis that these candidates would have greater translational relevance. UNCOVER was ran in greedy mode (UNCOVER_greedyv2.py) to identify negative association with a mutated gene sets of maximum 10 genes. To evaluate the confidence of association, we set the number of permutations as 100 to compute p-values and applied a threshold of p<0.01. Of the 633,232 predicted synthetic lethal candidates predicted by TCGA_DEPMAP_, 28,609 pairs also had evidence of mutually exclusive mutation rates in TCGA. The candidate list was then refined further by prioritizing paralogs using the biomaRt paralog database (version 2.28.0) R package. We additionally included pairs characterized by phylogenetic distance with threshold less than 1.5, as described previously (*71, 72*). The candidate list received a final filtering based on overall patient prevalence of LOF events, protein-protein interactions with TSG (*73, 74*), prior experimental evidence of gene-gene interactions (*6, 16, 17, 41, 42*), and manual curation to include essential and non-essential controls. The final list of gene pairs that were prioritized for experimental validation included 601 synthetic lethality candidates from the original lasso regression of TCGA_DEPMAP_ and an additional 264 pairs that were retained as library controls (**Table S8**).

### Multiplexed screening synthetic lethalities using AsCas12a (AsCpf1) and enAsCas12a (enAsCpf1)

Guides were designed using the TTTV PAM for AsCas12a and synthesized into 4-guide arrays with direct repeats (DR)-1, -2, -3, and -4 preceding each guide, followed by cloning into a guide-only lentiviral vector (pRDA_052), as described previously (*47, 48*). A double knockout construct was designed with 2 guides x 2 genes (n = 4 guides total per construct) for each pair of synthetic lethal candidates. Single knockout constructs were also designed 2 guides x 1 gene + 2 non-targeting guides (n= 4 guides total per construct) for each pair of synthetic lethal candidates. For some pairs, multiple single knockouts were used to assess overall library variance and were collapsed to the median values for downstream gene interaction analysis. A total of 500 constructs with 4 non-targeting guides were also included in the library as negative controls. An initial set of pilot screens were performed in triplicate using A549, NCI-H1299, MDA-MB-231, PC3M, and DETROIT562 that stably express AsCas12a, as described previously (*48*). An enhanced AsCas12a (enAsCas12a) enzyme was recently reported that is compatible with CRISPR/AsCas12a libraries (*46*), enabling an independent replication of the initial pilot screens and expansion to a total 14 total cancer cell models. The subsequent screens using enAsCas12a were performed in triplicate using A549, NCI-H1299, MDA-MB-231, NCI-H1703, PC3M, DETROIT562, HT29, HCT116, PANC1, MIAPACA2, SNU1, HSC2,

HSC3, and FADU. For all screens, cells were infected at a multiplicity of infection (MOI) = 0.3 and cultured for 14 days while continuously maintaining 500X coverage, followed by DNA extraction and PCR-barcoding using the p5 Agon and p7 Kermit primers (*48*). The PCR-barcoded libraries were single-end sequenced using an Illumina HiSeq4000 (300X cycle), followed by demultiplexing of sequencing reads (bcl2fastq, Illumina) and quantification of guide array abundance across all samples was done with a custom Perl script. Sequences between the flanking sequences or by location were extracted and compared to a database of sgRNA for each library. Only perfectly matched sgRNA sequences were kept and used in the generation of count matrix. Normalization between all samples was done using the “TMM” method (*75*) implemented in the edgeR R Bioconductor package. Log2 fold-changes (L2FC) of guide array abundance were calculated by comparing day 14 libraries with the plasmid library using limma-voom (*76*). Genetic interactions (GI) were calculated by comparing the expected and observed L2FC of double and single knockout constructs, as described previously (*41, 47*). Briefly, the expected L2FC for double knockout constructs is calculated as a sum (LF2C) of the individual knockout (sgRNA + Nontargeting). Synthetic lethal and buffering interactions are defined for double knockout in which the observed double knockout L2FC is significantly greater or less than that of the expected L2FC, respectively.

### Experimental validation of *PAPSS1/2* synthetic lethality

CRISPR/Cas12 knockouts of *PAPSS1* and/or *PAPSS2* were performed with Integrated DNA Technologies (IDT) Cas12 *Ultra* according to manufacturer’s instructions by Neon electroporation of RNPs (Invitrogen). Guides were designed using the Broad Institute CRISPick algorithm and the two best performing guides for each gene were used in combination; PAPSS1_guide#1 AGGATCGCAACAATGCAAGGCAA, PAPSS1_guide#2 TCTTCATGAGTCGCAGTCAGAAC, PAPSS2_guide#1 AGAACATTGTACCCTATACTATA, PAPSS2_guide#2 GAGGTGCATCTACAAATATTTCA. Protein expression was quantified by Simple Western (ProteinSimple, BioTechne) using the following antibodies; PAPSS1 clone 1F4 (Abnova, H00009061-M05) at 1:100, PAPSS2 (Cell Signaling Technology (CST), #70638) at 1:50, *PTEN* (CST, #9552) at 1:100 with beta-Actin clone 8H10D10 (CST, #3700) and Gapdh clone 14C10 (CST, #2118) as loading controls. Flow cytometry analysis of sulfonated HSPGs was performed with 10E4 antibody conjugated to FITC and used at 1:200 (USBiological Life Sciences, #H1890-10). Bacteroides Heparinase III was obtained from New England Biolabs (NEB, P0737L) and used as per manufacturer’s protocol by treating cells for 1hr in reaction buffer at 30°C before FACS analysis. Spheroid cultures were performed on Ultra-low attachment 96-well plates (Corning, #7007), growth was tracked on Incucyte S3 (Sartorius) and CellTiterGlo (CTG) readouts were performed for viability measurements (Promega, Cat# G9681). For rescue experiments, Heparan Sulfate (HS) was used at 10 to 50ug/mL (Sigma, H7640). For in vivo experiments, 1e6 UMUC3 cells were reconstituted in Hanks Balanced Salt Solution (HBSS), mixed 1:1 with Matrigel (Corning, 356235) and 200ul inoculated in the right flank (n=5 SCID/Beige mice per condition). Tumors were extracted at day 22, mechanically dissociated with scalpels and single-cell suspensions were made using Liberase and DNAseI (Millipore Sigma, Cat 05401127001 and 11284932001) incubated at 37oC for 1hr and mouse cells were magnetically depleted on LS columns using mouse cell depletion cocktail (Miltenyi, 130-104-694 and 130-042-401).

### Characterization of PDXE_DEPMAP_

The distribution of the cross-validated expression-only models of gene essentiality (n=1,996) across lineages was assessed by unsupervised cluster analysis (Ward.D2 method) and visualized using the ComplexHeatmap R package (version 2.6.2). Associations of dependencies with genomic features were assess using a Wilcoxon Rank Sum differential test as implemented using stat_compare_means function of ggpubr R package (version 0.4.0). To test the ability of gene essentiality to predict the response to corresponding targeted therapies, the change in PDX burden from baseline to experimental endpoint was correlated with target gene essentiality in PDXE_DEPMAP_ using a Pearson’s correlation test and FDR correction of p-values for multiple hypothesis testing. Receiver operating characteristic (ROC)–area under the curve (AUC) analysis was performed using the pROC R package (version 1.18.0) to assess the accuracy of drug responses predicted by the target gene essentiality scores. Only drugs with at least 20 treated PDX models were evaluated, and the metrics are reported in **Table 1**.

### Characterization of GTEX_DEPMAP_

The distribution of the cross-validated expression-only models of gene essentiality (n=1,996) across normal tissues was assessed by unsupervised cluster analysis (Ward.D2 method) and visualized using the ComplexHeatmap R package (version 2.6.2). Differences in gene essentiality in normal and malignant tissues, as well as malignant tissues with genomic features, were assessed using a Wilcoxon Rank Sum differential test as implemented using stat_compare_means function of ggpubr R package (version 0.4.0). To evaluate the sensitivity and specificity of GTEX_DEPMAP_ to genes associated with tissue-specific toxicities, we profiled GTEX_DEPMAP_ genes associated with both blood disorders and drug-induced liver toxicity using the Cortellis OFF-X database (https://targetsafety.info/). The OFF-X database is a drug and target safety intelligence database that predicts potential associations based off both preclinical and clinical safety data alerts from peer-reviewed journals, company communications, clinical trials, and regulatory agency communications. These blood and liver toxicity associations were further evaluated to identify overlapping or unique genes to each toxicity and annotated with the frequency of associated safety alerts. In total, the Cortellis OFF-X database identified for drug targets associated with potential toxicities in blood (n=82), liver (n=85), or blood and liver (n=74), which were then compared across normal tissue lineages in GTEX_DEPMAP_. To compare gene essentiality between malignant and normal tissues, TCGA_DEPMAP_ and GTEX_DEPMAP_ samples were matched based on the tissue-of-origin and a student’s t-test was applied to differential analysis between the dependency profiles of tumor and normal tissue of the same lineages. The t-statistic was used to characterize the dependency difference between the tumor and corresponding normal tissue with a negative t-statistic value corresponding to a higher dependency in tumor as compared with the normal tissue. Gene ontology enrichment analysis (GSEA) was performed across all paired malignant and normal tissue-of-origin. The list of genes for the lung network was generated using the top 100 genes showing the largest differentiation in gene essentiality between cancer compared to normal tissue in lung based on the negative t-statistic values. Network connectivity and the gene ontology enrichment were calculated using STRING (https://string-db.org/), as described previously (*77*).

