## Supplemental Figures for "TCGA_DEPMAP_ – Mapping Translational Dependencies and Synthetic Lethalities within The Cancer Genome Atlas"

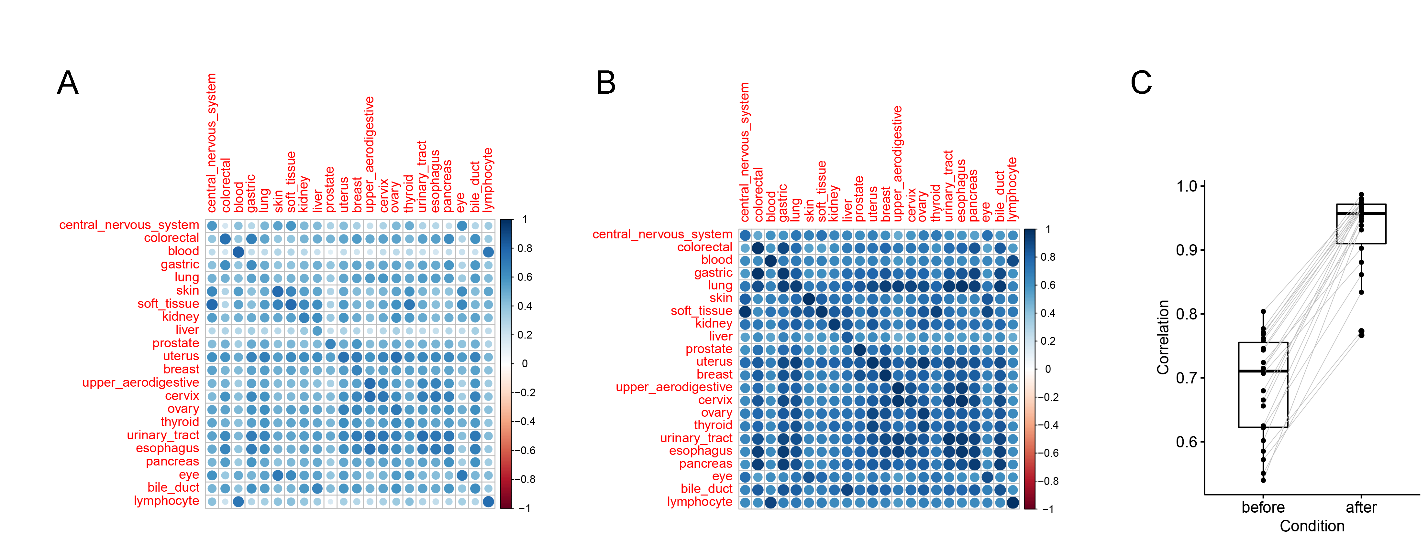

**Figure S1.** Model performance is significantly improved by identification and removal of the most variant signatures (cPC1-4; i.e., stromal signatures) before elastic net ML. The heatmaps show the Pearson correlation between the gene expression of DepMap and TCGA before (A) and after (B) expression alignment, where rows are TCGA lineages and columns are DepMap lineages. It shows that the correlation of expression for the same lineage in TCGA and DepMap is significantly improved by our expression alignment pipeline (C).

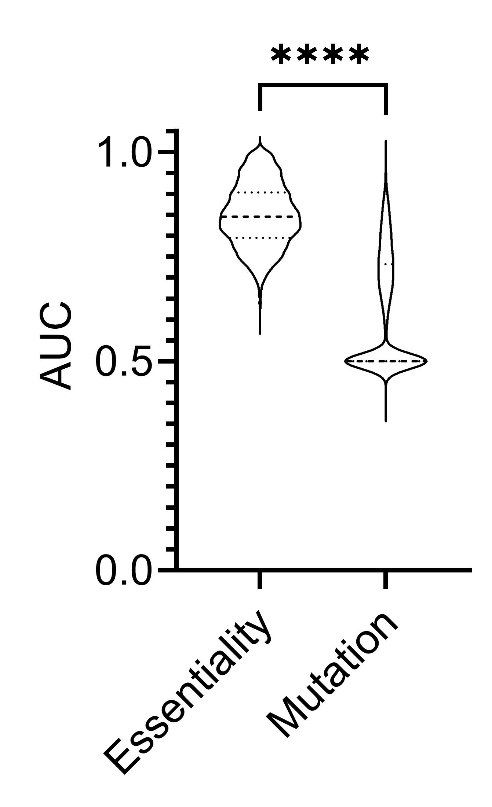

**Figure S2.** Comparison of expression-only elastic-net models for gene essentiality and gene mutational status. To make performance metrics (AUC) comparable with binary mutational status, the essentiality scores were binarized using a –0.5 essentiality score as a cutoff. To calculate the accuracy of predicting dependencies and mutations, elastic-net machine learning was run to predict mutations and essentiality using the same settings and expression data for 891 genes with mutations at >2% prevalence in TCGA_DEPMAP_ patients. Of note, the elastic-net models were allowed to select the most informative predictive features for mutation and essentiality for each gene, as the best predictors for essentiality may not be the best features to predict mutation. *****P*<0.0001 by Student unpaired t-test.

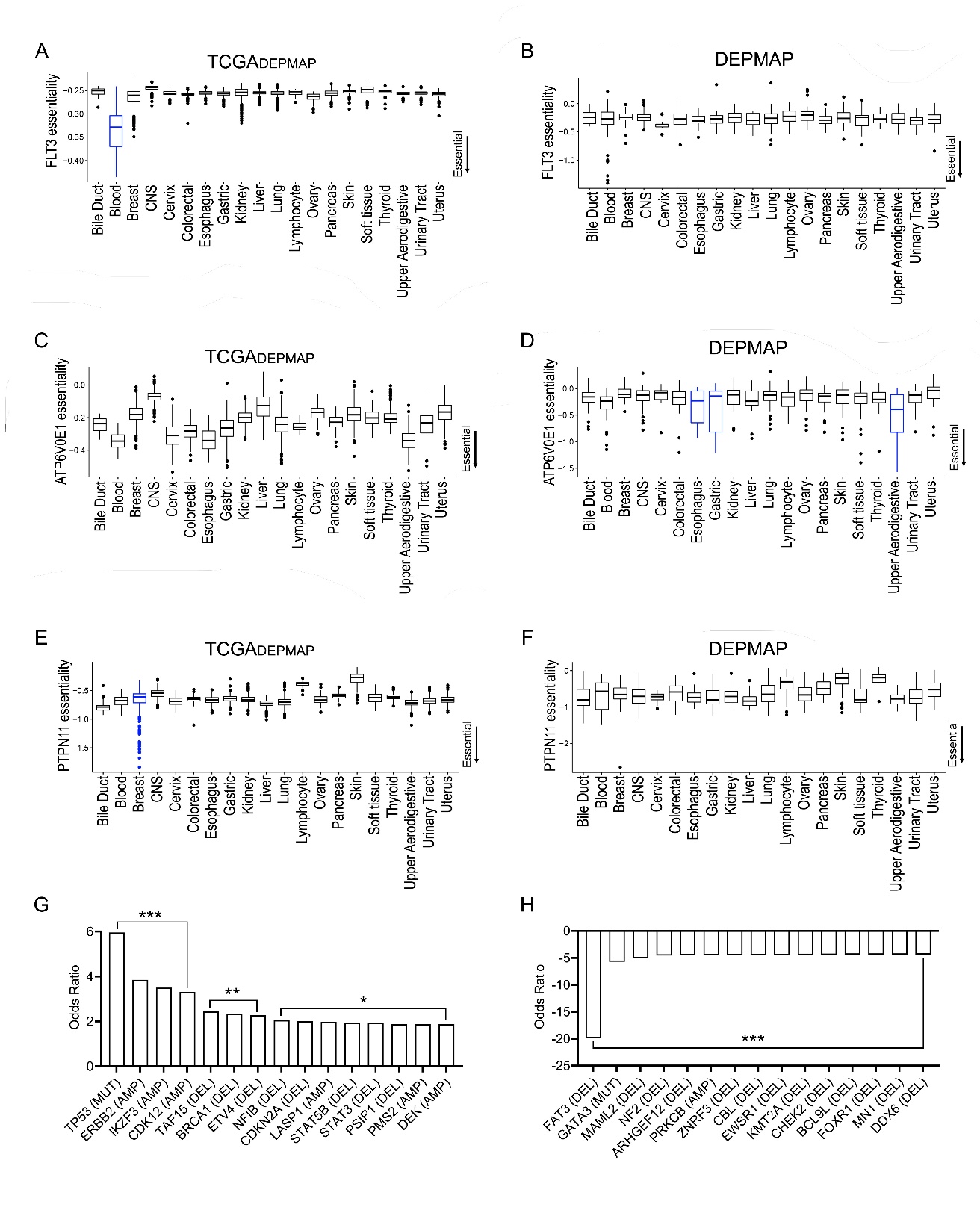
**Figure S3.** Examples of dependencies with different selectivity profiles across TCGA_DEPMAP_ and DEPMAP cohorts. (**A**) *FLT3* was classified as a strongly selective dependency (SSD) with markedly higher dependency in blood lineage cancers of TCGA_DEPMAP_ (blue bar), (**B**) whereas *FLT3* showed higher dependency in some blood lineage cancers but does not meet the threshold of an SSD in DEPMAP. (**C**) *ATPV6V0E1* essentiality scores varied widely across TCGA_DEPMAP_, (**D**) while *ATPV6V0E1* was classified as an SSD that was restricted to only a few lineages in DEPMAP (blue bars). (**E**) *PTPN11* was classified as an SSD with very strong dependencies in a subset of breast cancer patients in TCGA_DEPMAP_ (blue bar), (**F**) whereas no selectivity of *PTPN11* essentiality was detected in DEPMAP. (**G**) Top cancer driver mutations enriched in TCGA_DEPMAP_ breast cancer patients that were highly dependent on *PTPN11*. (**H**) Top cancer driver mutations depleted in TCGA_DEPMAP_ breast cancer patients that were highly dependent on *PTPN11*. For (**G**, **H**), ***FDR<0.01, ***P*<0.01, and **P*<0.05, as determined by Fisher exact test.

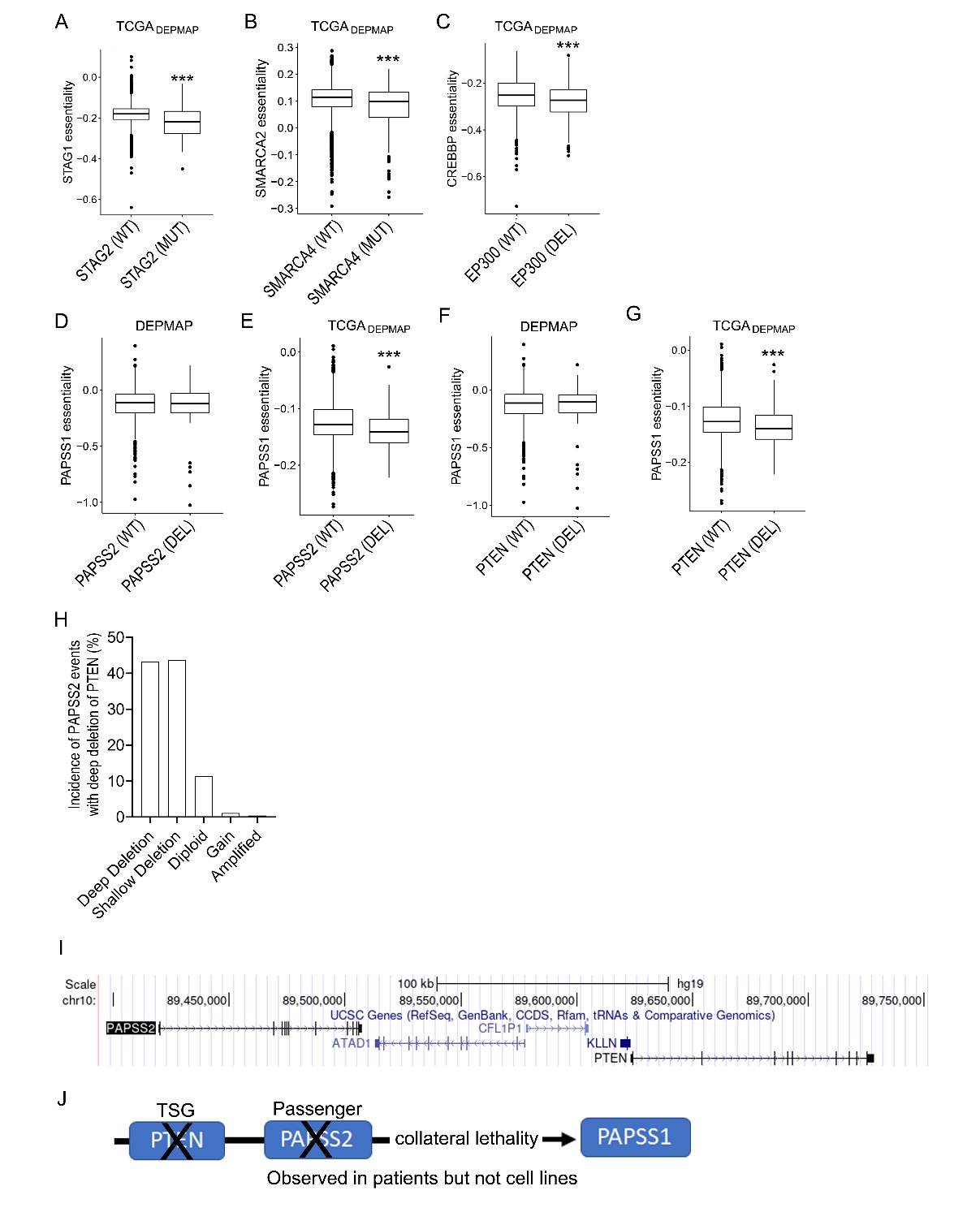
**Figure S4.** Examples of synthetic lethalities detected by TCGA_DEPMAP_. (**A**) *STAG1* synthetic lethality with *STAG2* mutation, (**B**) *SMARCA2* synthetic lethality with *SMARCA4* mutation, and (**C**) *CREBBP* synthetic lethality with *EP300* mutation are examples of well-known synthetic lethalities that were detected by TCGA_DEPMAP_. (**D**, **E)** *PAPSS1* is a novel synthetic lethality in the context of *PAPSS2* deletion, which is not detectable in (**D**) DEPMAP cell lines and is only detectable in (**E**) TCGA_DEPMAP_ patient samples. (**F**, **G**) Likewise, *PAPSS1* is not synthetic lethal with *PTEN* deletion in DEPMAP cell lines (**F**) and is only detectable in TCGA_DEPMAP_ patient samples (**G**). (**H**) Unlike cultured cell models, *PAPSS2* is frequently co-deleted with *PTEN* in TCGA patients. (**I**) *PAPSS2* is a closely neighboring gene of *PTEN*. (**J**) A schematic representation summarizing the hypothesized synthetic lethality of *PAPSS1* that is driven by collateral deletion of *PAPSS2* with the tumor suppressor gene (TSG), *PTEN*, in patients but not cell lines. ****P* < 0.001, as determined by the Wilcoxon rank sum test.

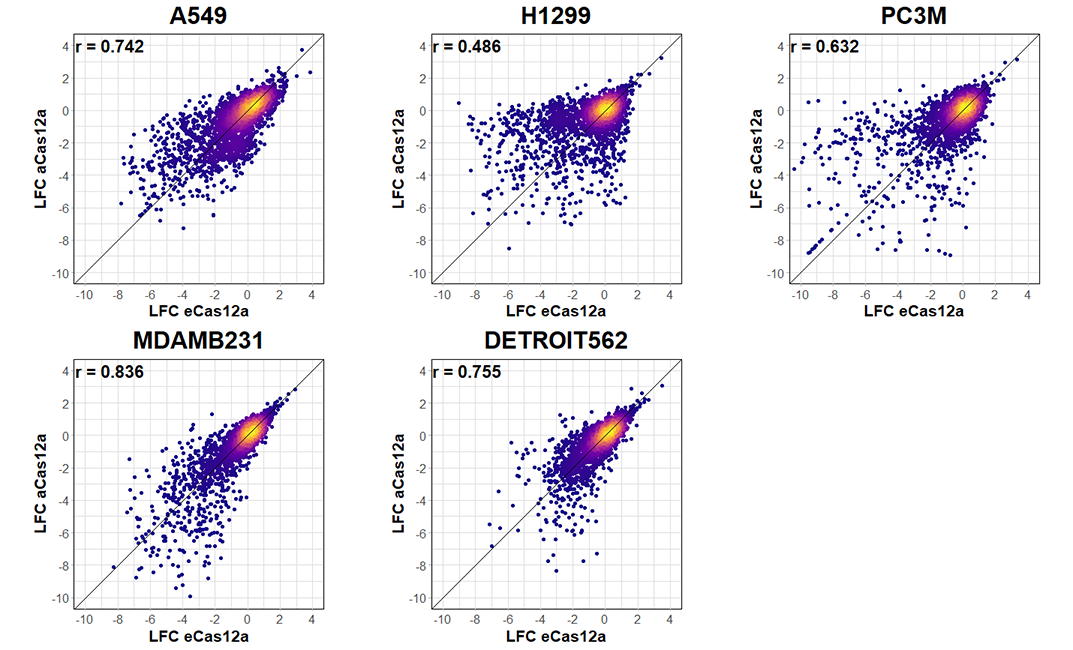

**Figure S5.** Comparison of multiplexed CRISPR/Cas12 screens performed using AsCas12a and EnAsCas12a enzymes. Analysis was performed using a Pearson’s correlation and coefficients (*r*) are displayed on the graphs.

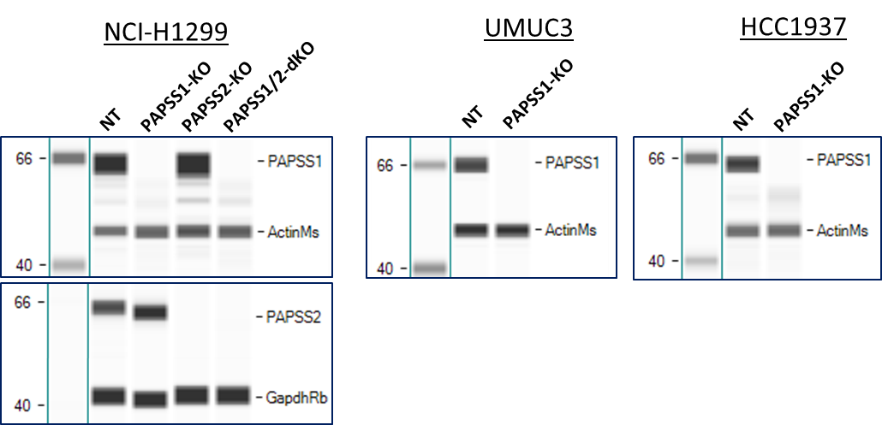

A

B

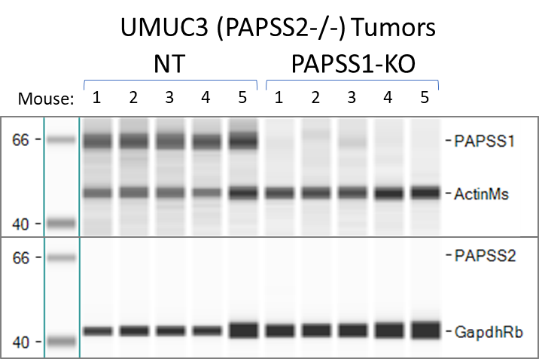

C

NCI-H1299

UMUC3

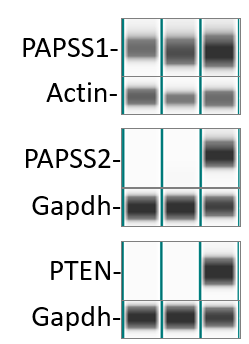

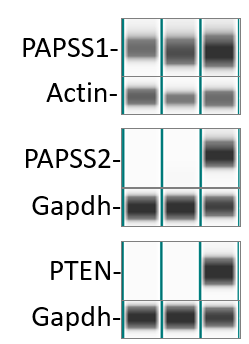

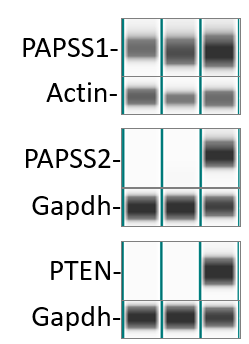

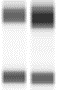

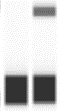

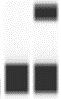

**Figure S6.** Simple western analysis of PAPSS1, PAPSS2, PTEN and loading controls from experimental assays. (**A**) Endogenous expression of PAPSS1, PAPSS2, and PTEN in the model cell lines UMUC3 and NCI-H1299. (**B**) Validation of PAPSS1 and PAPSS2 single (KO) and double (dKO) knockouts by RNP in spheroid experiments. (**C**) Validation of PAPSS1 knockout in the UMUC3 xenograft experiment tumors.

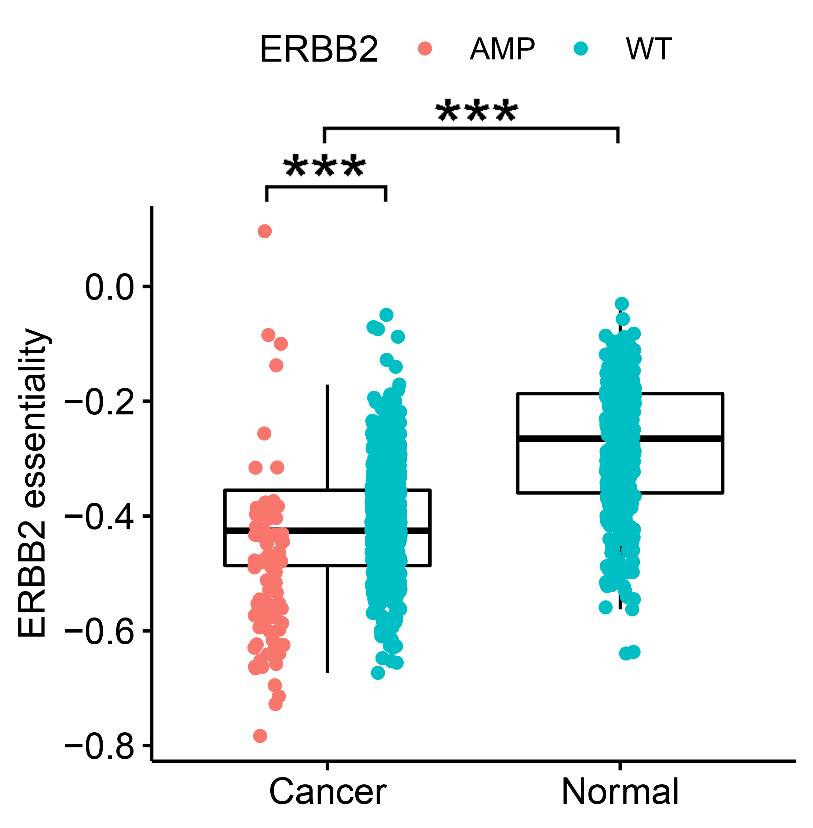

**Figure S7.** ERBB2 essentiality is significantly higher in malignant breast cancer with ERBB2 amplifications (TCGA_DEPMAP_) compared with normal breast (GTEX_DEPMAP_). ****P* < 0.001, as determined by the Wilcoxon rank sum test.

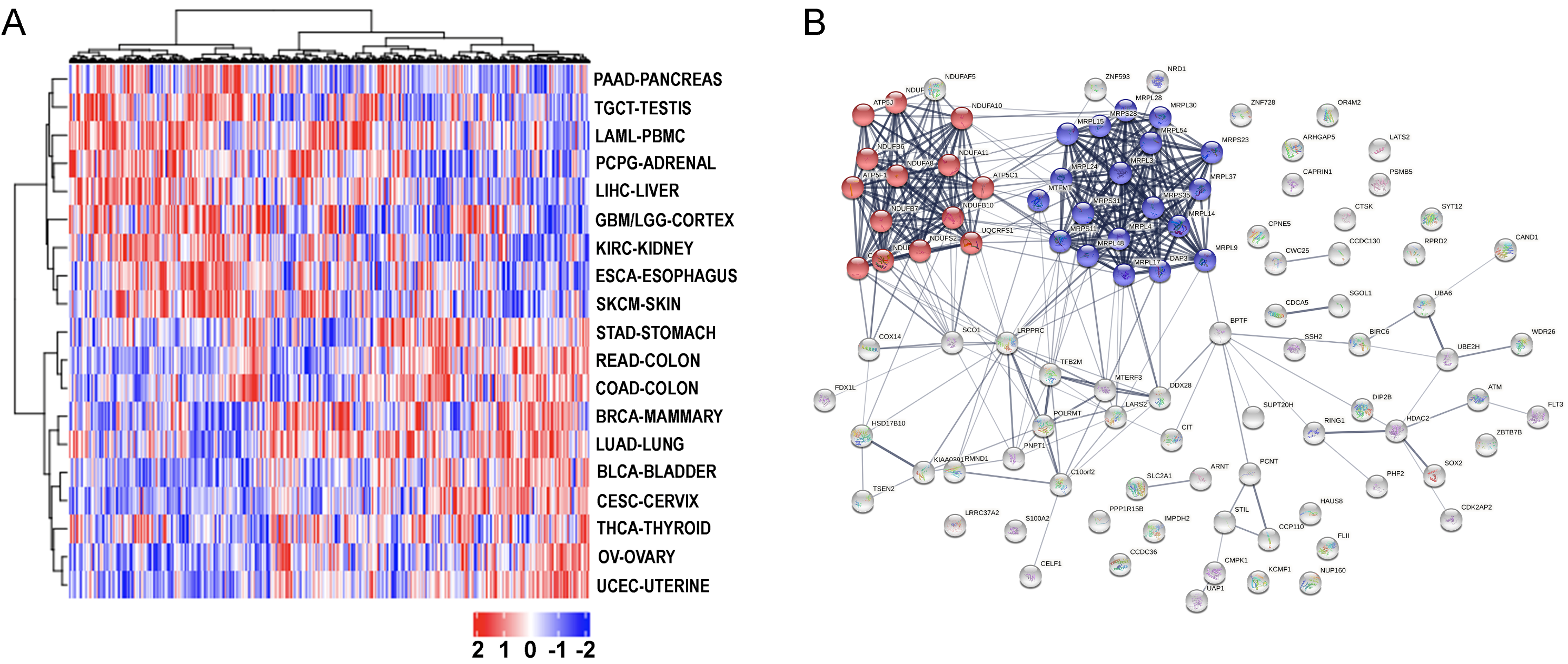

**Figure S8.** STRING network analysis of the top 100 LUAD targets with the greatest predicted tolerability in normal lung reveals significant connectivity (p<1x10^-16^) and gene ontology enrichment for oxidative phosphorylation (blue colored spheres; p=5.8x10^-11^) and mitochondrial translation (red colored spheres; p=2.9x10^-20^).

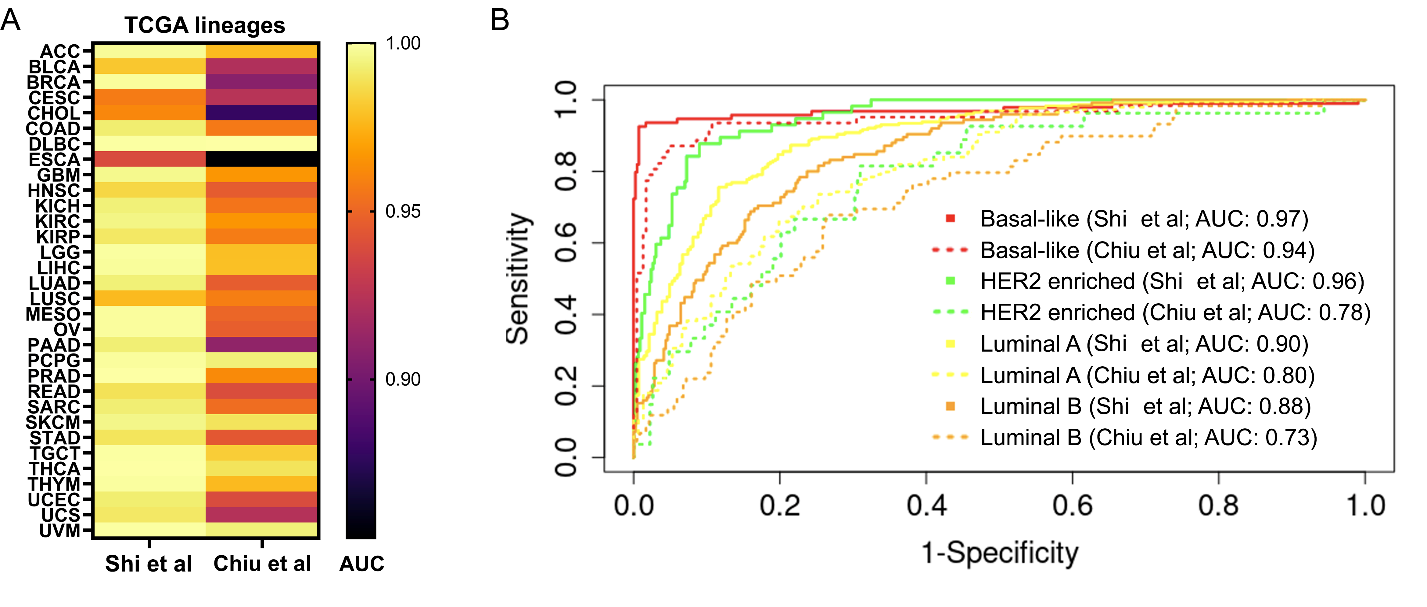

**Figure S9.** TCGA_DEPMAP_ outperforms DeepDEP. (A) Precision-recall analysis of pancancer lineage predictions by the AUC values are significantly higher for TCGA_DEPMAP_ in predicting cancer lineages based on top 100 variable dependencies compared with DeepDEP. (B) The ROC curves for predicting the breast cancer subtypes based on the top 100 variable gene dependencies. The TCGA_DEPMAP_ significantly outperforms DeepDEP in predicting any of the breast cancer subtypes (TCGA_DEPMAP_ continuous line; DeepDEP dotted line).
