## Supplemental Table Legend for "TCGA_DEPMAP_ – Mapping Translational Dependencies and Synthetic Lethalities within The Cancer Genome Atlas"

Table S1. Expression-only elastic-net model coefficients of gene essentiality predicted from the DEPMAP. The rows denote features used in modeling and columns are corresponding gene models.

Table S2. Multi-omics elastic net model coefficients of gene essentiality predicted from the DEPMAP. The rows denote features used in modeling and columns are corresponding gene models.

Table S3. List of cross-validated genes from expression-only and multi-omics models.

Table S4. Pathway enrichment analysis of dependencies most changed by the transcriptional alignment, as defined by gene essentiality scores with the least correlation (r < 0.5) before and after alignment of DEPMAP and TCGA transcriptomes.

Table S5. Gene essentiality scores across TCGA_DEPMAP_. The row labels denote the gene model and the column labels correspond to TCGA patient sample ID. ([Table S5 link](https://figshare.com/articles/dataset/Supplemental_Tables_Final/21558834))

Table S6. Strongly selective dependencies (SSDs) across DEPMAP and TCGA_DEPMAP_, as assessed by the normality likelihood ratio test (NormLRT).

Table S7. Cox log rank analysis of gene essentiality associated with progression free interval (PFI) in TCGA_DEPMAP_. The row labels denote the gene models and the column heads correspond to the hazard ratios (HR) and p-values (PV), respectively.

Table S8. Annotation of synthetic lethality candidates that were experimentally tested by multiplexed CRISPR/Cas12 screening. The row labels denote gene pairs and column labels correspond to whether a pair was predicted by TCGA_DEPMAP_ and whether the pair was annotated as paralogs or tumor suppressor genes (TSG).

Table S9. Gene interaction scores from CRISPR/Cas12 screens using AsCas12a and enAsCas12a across 14 cell models. The row labels denote the gene pairs and the column labels correspond the to the enzyme used (AsCas12a or enAsCas12a), the expected (exp.) effect (summed z-score of the single knockouts), the observed (obs.) effect of double knockout (z-score transformed), the difference (Diff.) of expected and observed effects, and the z-score transformation of the difference (Diff_Z).

Table S10. Gene essentiality scores across PDXE_DEPMAP_. The row labels denote the gene model and the column labels correspond to the PDX sample ID. ([Table S10](https://figshare.com/articles/dataset/Supplemental_Tables_Final/21558834))

Table S11. Gene essentiality scores across GTEX_DEPMAP_. The row labels denote the gene model and the column labels correspond to the GTEX sample ID. ([Table S11](https://figshare.com/articles/dataset/Supplemental_Tables_Final/21558834))

Table S12. List of genes associated with both blood disorders and drug-induced liver toxicity that were curated from the Cortellis OFF-X database. The OFF-X database is a drug and target safety intelligence database that predicts potential associations based off both preclinical and clinical safety data alerts from peer-reviewed journals, company communications, clinical trial, regulatory agency communications. The rows denote the genes associated with potential toxicities in blood (n=82), liver (n=85), or blood and liver (n=74). The column labels correspond to the different evidence classes of safety alerts.

Table S13. Differences between predicted essentiality between malignant tissue and the normal tissue-of-origin. The rows denote the gene model and the column labels correspond to the malignant (TCGA_DEPMAP_) and normal (GTEX_DEPMAP_) tissue types that were matched based on the tissue-of-origin. A Student’s t-test was applied to differential analysis between the dependency profiles of tumor and normal tissue of the same lineages. The t-statistic was used to characterize the dependency difference between the tumor and corresponding normal tissue with a negative t-statistic value corresponding to a higher dependency in tumor as compared with the normal tissue.

Table S14. Pathway analysis of the strongest tumor dependencies with the least normal tissue-of-origin toxicity. The rows denote the pathway and the column labels correspond to the enrichment p-value (pval), FDR-corrected p-value (padj), normalized enrichment score (NES), number of genes included in the gene set (size), the gene models comprising the leading edge (leadingEdge), and the paired malignant and normal tissues (type).
